# Prostate cancer resistance leads to a global deregulation of translation factors and unconventional translation of long non-coding RNAs

**DOI:** 10.1101/2021.01.05.425492

**Authors:** Emeline I. J. Lelong, Pauline Adjibade, France-Hélène Joncas, Gabriel Khelifi, Valerie ST.-Sauveur Grenier, Amina Zoubedi, Jean-Philippe Lambert, Paul Toren, Rachid Mazroui, Samer M. I. Hussein

## Abstract

Emerging evidence associates translation factors and regulators to tumorigenesis. Recent advances in our ability to perform global translatome analyses indicate that our understanding of translational changes in cancer resistance is still limited. Here, we generated an enzalutamide-resistant prostate cancer (PCa) model, which recapitulated key features of clinical enzalutamide-resistant PCa. Using this model and polysome profiling, we investigated global translation changes that occur during the acquisition of PCa resistance. We found that enzalutamide-resistant cells exhibit a discordance in biological pathways affected in their translatome relative to their transcriptome, a deregulation of proteins involved in translation, and an overall decrease in translational efficiency. We also show that genomic alterations in proteins with high translational efficiency in enzalutamide-resistant cells are good predictors of poor patient prognosis. Additionally, long non-coding RNAs in enzalutamide-resistant cells show increased association with ribosomes, higher translation efficiency, and an even stronger correlation with poor patient prognosis. Taken together, this suggests that aberrant translation of coding and non-coding genes are strong indicators of PCa enzalutamide-resistance. Our findings thus point towards novel therapeutic avenues that may target enzalutamide resistant PCa.

## INTRODUCTION

Translation is one of the last processes in the flow of genetic information. It is a multistep and highly controlled protein synthesis process consisting of three major steps, namely initiation, elongation and termination (Hershey et al. 2019). Translation initiation depends on a network of interacting translation initiation factors (eIFs) which are highly regulated. In particular, regulation of the activity and expression of two eIFs: eIF4F and eIF2A, is under extensive study, revealing an important role for translation regulation in cellular processes such as cell differentiation, growth and cell stress response (Holcik and Sonenberg 2005; Chang and Stanford 2008). Dysregulation of eIFs can be linked to various cancers. Altered expression or activity of components of the eIF4F complex such as *EIF4E* and *EIF4G*, have been observed to support cancer cell growth by activating translation initiation of mRNAs encoding key cell cycle regulators, as well as survival and oncogenic factors (Bhat et al. 2015). Furthermore, *EIF4E* phosphorylation promotes prostate tumorigenesis and is elevated in castrate resistant prostate cancer (PCa). This correlates with disease progression and poor clinical outcomes in patients with PCa (Furic et al. 2010). On the other hand, while phosphorylation of *eIF2A* blocks general translation in cases of cellular stress, it allows the preferential translation of a specific set of target mRNAs involved in cell adaptation to stress and survival (Pakos‐Zebrucka et al. 2016). This is in line with recent evidence associating alterations in the phosphorylated *EIF2A* translational pathway with cancer, a process highly linked to the cellular stress response (Koshikawa et al. 2006). Moreover, altered phosphorylation of *EIF2A* has been observed to occur as an adaptive stress response in both murine and humanized models of aggressive and resistant PCa (Nguyen et al. 2018). Perturbations in translation regulation may therefore represent key indicators of PCa severity.

PCa resistance is a highly prevalent and common cause of cancer-related death worldwide (Bray et al. 2018; Siegel et al. 2020). Despite effective local treatments, many patients experience recurrences and eventually develop metastases (Vickers et al. 2008; Boorjian et al. 2012; KRYGIEL et al. 2005). Highly dependent on androgens for growth, recurrent or metastatic PCa is treated with androgen deprivation therapy (ADT). Concomitant or subsequent use of enzalutamide (ENZ), a potent androgen receptor (AR) antagonist, significantly delays the consequences of treatment failure (Beer et al. 2014; Armstrong et al. 2019; Davis et al. 2019). However, not all patients benefit from the therapeutic effects of ENZ and all eventually develop resistance (Buttigliero et al. 2015). This highlights an urgent need to find reliable markers that can predict patient response and development of resistance. Recent genomics studies have led to the discovery of promising PCa biomarkers (Mikropoulos et al. 2014; Ngollo et al. 2014; Peng et al. 2014). Due to the relative ease of nucleic acid sequencing, a large majority of existing PCa-related data focuses on transcriptomic studies analysing total RNA abundance as a stand-in for protein levels. This precludes discovery of many potential biomarkers whose protein expression relies mainly on the translational rate. Indeed, it is now well established that transcriptomic estimates of RNA abundance alone are insufficient to capture proteins whose differential expression critically impact cellular differentiation and growth, environmental and pathological stress, or tumorigenesis (Schwanhäusser et al. 2011; Maier et al. 2009). This is in part due to the complex regulatory mechanisms that orchestrate the translation of RNAs. It is estimated that about 40 % of protein level variations are due to translational regulation. Thus, accurate estimation and identification of relevant protein variations occurring in various cancers including PCa calls for integrative methods that measure the transcriptome, the RNAs associated with translating poly(ribo)somes (translatome), as well as the proteome.

Monitoring the translational status of entire transcripts via polysome profiling and RNA sequencing (RNA-seq) is a powerful approach used to identify ribosome-associated RNAs (Coudert et al. 2014). Indeed, several polysome profiling studies on cancer cell lines have recently succeeded to identify cancer cell-specific signatures not detected by standard RNA-seq analyses (Lupinacci et al. 2019; Kusnadi et al. 2020; Wahba et al. 2016). Hsieh *et al.* reported the first study in PCa using polysome profiling (Hsieh et al. 2012); however, they analysed AR-negative PCa cells in their work, which may not be a direct reflection of the acquisition of ENZ-resistance observed in patients. They found that *eIF4F*, driven by its upstream mTORC1 signaling regulatory pathway, promotes a metastatic phenotype in PCa through preferential translation of mRNAs encoding proteins involved in cell invasion and metastases. This is consistent with data revealing the PI3K/AKT/mTOR translational pathway as a key oncogenic pathway in treating resistant PCa (TOREN and ZOUBEIDI 2014). Even though accumulating evidence supports a potential role played by translation regulation in the progression of PCa, the role of translational changes in the acquisition of ADT resistance or ENZ-resistance in PCa remains unknown.

To investigate perturbations in the translatome acquired upon PCa ENZ-resistance, we utilize an integrative approach merging global analysis of RNAs by RNA-Seq and of their association to ribosomes through poly(ribo)some profiling in ENZ-sensitive and ENZ-resistant PCa cell lines. We apply this method to a novel model of castration-resistant (ENZ-sensitive) and ENZ-resistant PCa we developed from the well-known AR-positive VCaP prostate cancer cell line (Korenchuk et al. 2001). These analyses are complemented by use of the ENZ-resistant MR49F cell line and its sensitive parental cell line LNCaP (Bishop et al. 2017). These results were corroborated by mass spectrometry and publicly available gene expression data, suggesting that translation is indeed globally altered during acquisition of resistance. Furthermore, our analysis revealed enrichment of long non-coding RNAs (lncRNAs) associated to ribosomes, which may suggest aberrant translation of novel peptides in the context of ENZ-resistant PCa. Our findings thus point towards novel biomarkers or therapeutic targets which are involved in PCa resistance to treatment.

## RESULTS

### Recapitulation and characterization of ENZ-resistant PCa

With the advent of potent AR-antagonists such as enzalutamide as first line therapy for castration-resistant prostate cancer (CRPC) patients, a few ENZ-resistant cellular models were developed (Simon et al. 2021; Hoefer et al. 2016). Among the first and most widely characterized ENZ-resistant cells (Kuruma et al. 2013; Toren et al. 2015; Bishop et al. 2017) were MR49F, generated through serial passage of LNCaP cells (androgen-sensitive prostate adenocarcinoma cells) in ENZ-treated mice (Kuruma et al. 2013). In an effort to develop a complementary model from human PCa cells with a wild-type AR (AR in LNCaP is mutated (Veldscholte et al. 1992)) (Table 1) and a concomitantly passaged castration-resistant control, we used a similar approach with the VCaP cell line (Kuruma et al. 2013). VCaP cells were inoculated in male athymic nude mice (Fig. 1A); mice were surgically castrated, and cells termed VCaP^CRPC^ were derived from tumors resistant to castration. In parallel, castrate-resistant tumors were treated with ENZ until regrowth, at which time the VCaP^ER^ cell line was established.

**Figure 1.**
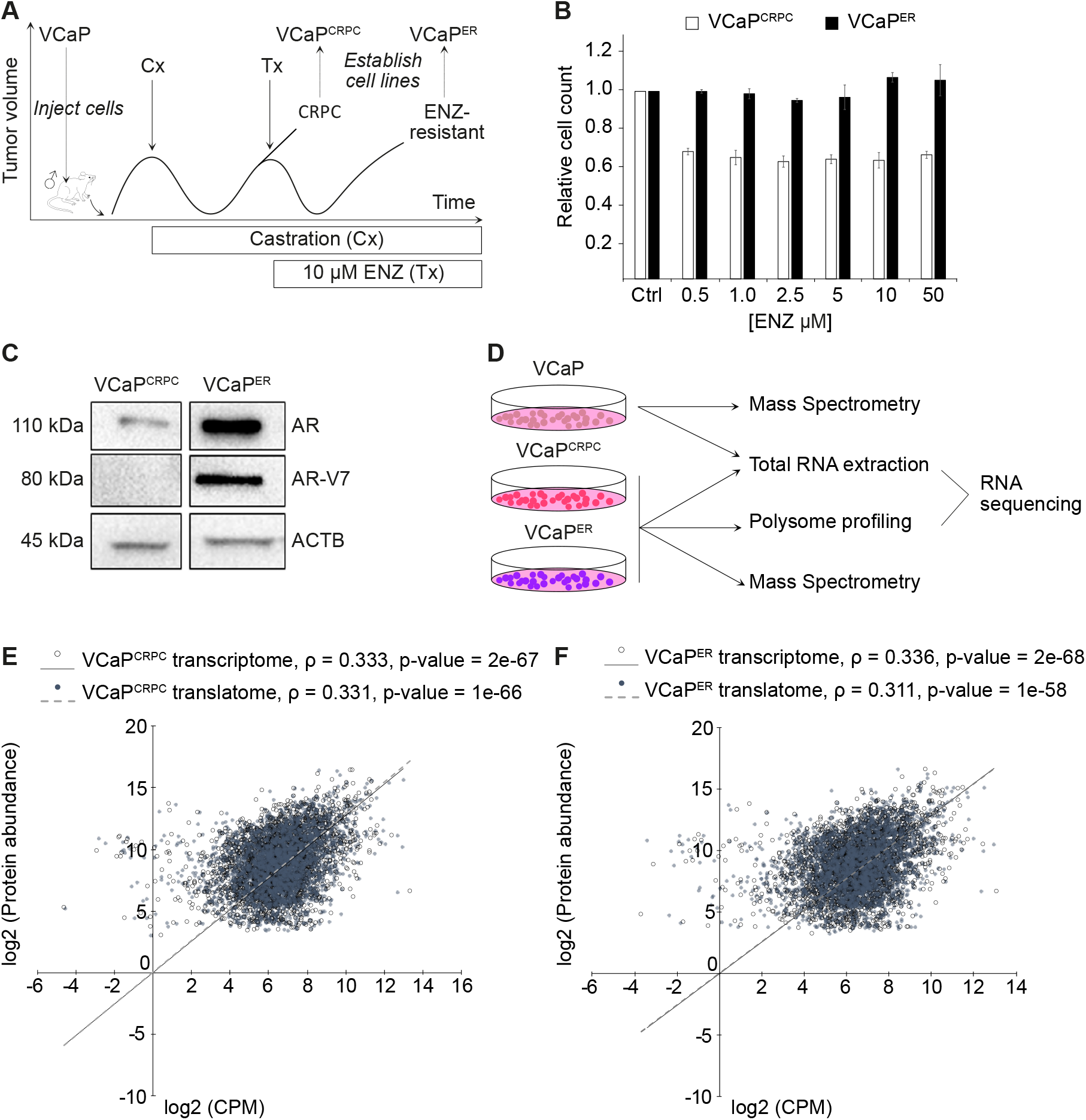
Establishment of Enzalutamide (ENZ)-resistant Prostate Cancer (PCa) cellular models. **(A)** Experimental approach to establish PCa resistance in mice model and to derive ENZ-sensitive and ENZ-resistant cell lines. Cx : surgical castration, Tx : ENZ treatment. **(B)** Cell proliferation assays on VCaP^CRPC^ and VCaP^ER^ performed with increasing quantities of ENZ. **(C)** Western-Blot showing expression of Androgen Receptor (AR) and resistance-specific splice variant AR-V7 in VCaP^ER^ and VCaP^CRPC^. **(D)** Analysis of transcriptome, translatome and proteome in PCa cell lines: Transcriptome and proteome from VCaP, VCaP^CRPC^ and VCaP^ER^ and translatome from VCaP^CRPC^ and VCaP^ER^. **(E)** Scatterplot and Pearson correlation analysis showing correlation between transcriptome and proteome (empty dots) and between translatome and proteome (full dots) of the VCaP^CRPC^ **(F)** and the VCaP^ER^ cell lines. Correlation coefficients (ρ) and linear regression (grey lines) are indicated.

To confirm acquisition of ENZ-resistance in VCaP^ER^ cells, we performed proliferation assays for VCaP^CRPC^ and VCaP^ER^ in the presence or absence of ENZ. ENZ treatment reduced cell proliferation of VCaP^CRPC^ cells while VCaP^ER^ cells were unaffected (Fig. 1B). Furthermore, we found that VCaP^ER^ displayed both increased AR and AR-V7 splice variant (Antonarakis et al. 2014) expression (Fig. 1C; Table 1). This variant encodes a truncated protein that lacks the C-terminal ligand-binding domain but retains the N-terminal domain and could therefore constitutively activate downstream target genes involved in PCa progression (Nadiminty et al. 2013; Mostaghel et al. 2011). AR-V7 expression is higher in advanced PCa and has been linked to ENZ resistance. These results indicate that our ENZ-resistant VCaP^ER^ cell line recapitulates key characteristics of clinical ENZ-resistant prostate cancer.

**Table 1.** Characteristics of AR in PCa cell lines.

| Cell line | Androgen Receptor | AR-V7 splice variant levels |
| --- | --- | --- |
| VCaP | Wild-type | Low-moderate |
| VCaP<sup>CRPC</sup> | Wild-type | Low-moderate |
| VCaP<sup>ER</sup> | Wild-type* | High |
| LNCaP | T878A mutant | Very low |
| MR49F | T878A and F876L mutant | Very low |

To understand how transcription and translation are coordinated during acquisition of ENZ-resistance in PCa, we analyzed the transcriptome (i.e. total RNA-seq), translatome (i.e. Heavy polysome-bound RNA-seq) and proteome (i.e. total proteins quantified by mass spectrometry (MS)) from the VCaP^CRPC^ and VCaP^ER^ cell lines, and the transcriptome and proteome from the parental VCaP cell line (Fig. 1D; Supplemental Table S1, Supplemental Table S2). Comparison of polysome profiles revealed no obvious differences between VCaP^CRPC^ and VCaP^ER^ (Supplemental Fig S1A, B). Additionally, through Pearson correlation coefficient (ρ) analysis, we found that the transcriptome and translatome correlated well in both cell lines (Supplemental Fig. S1C, D). We further observed relatively moderate correlations between transcriptome and proteome in VCaP^CRPC^ and VCaP^ER^ (ρ = 0.333 and ρ = 0.336 respectively) and between their translatome and proteome (ρ = 0.331 and ρ = 0.311 respectively) (Fig. 1E, F). Interestingly, although not significant, we do observe a modestly lower correlation between the translatome and proteome of the ENZ-resistant of VCaP^ER^ cell line (ρ = 0.311) relative to the other comparisons. This may indicate a slight shift in the translational landscape during prostate cancer ENZ-resistance.

### ENZ-resistance is accompanied by changes in the translatome

To provide a global view of changes to the translatome that are related to ENZ resistance, we performed differential expression analysis on total and heavy polysome-bound RNA-seq. We found several RNAs differentially bound to ribosomes between VCaP^ER^ and VCaP^CRPC^ (695 and 794, respectively) but not differentially expressed (Fig. 2A, B). Meanwhile, only 489 and 225 RNAs were upregulated in VCaP^ER^ and VCaP^CRPC^ respectively. Upon a GO term analysis, we observed similar GO term enrichment between the translatome and the transcriptome of VCaP^CRPC^, with several processes linked to cell membrane, cell adhesion, and development (Fig. 2C; Supplemental Fig. S2A; Supplemental Table S3). In contrast, the translatome and transcriptome of VCaP^ER^ showed different enriched GO terms. Interestingly, the VCaP^ER^ transcriptome showed enrichment for GO terms similar to the ones in VCaP^CRPC^, while the VCaP^ER^ translatome GO terms were associated with transcription regulation and proteins localized to the nucleus.

**Figure 2.**
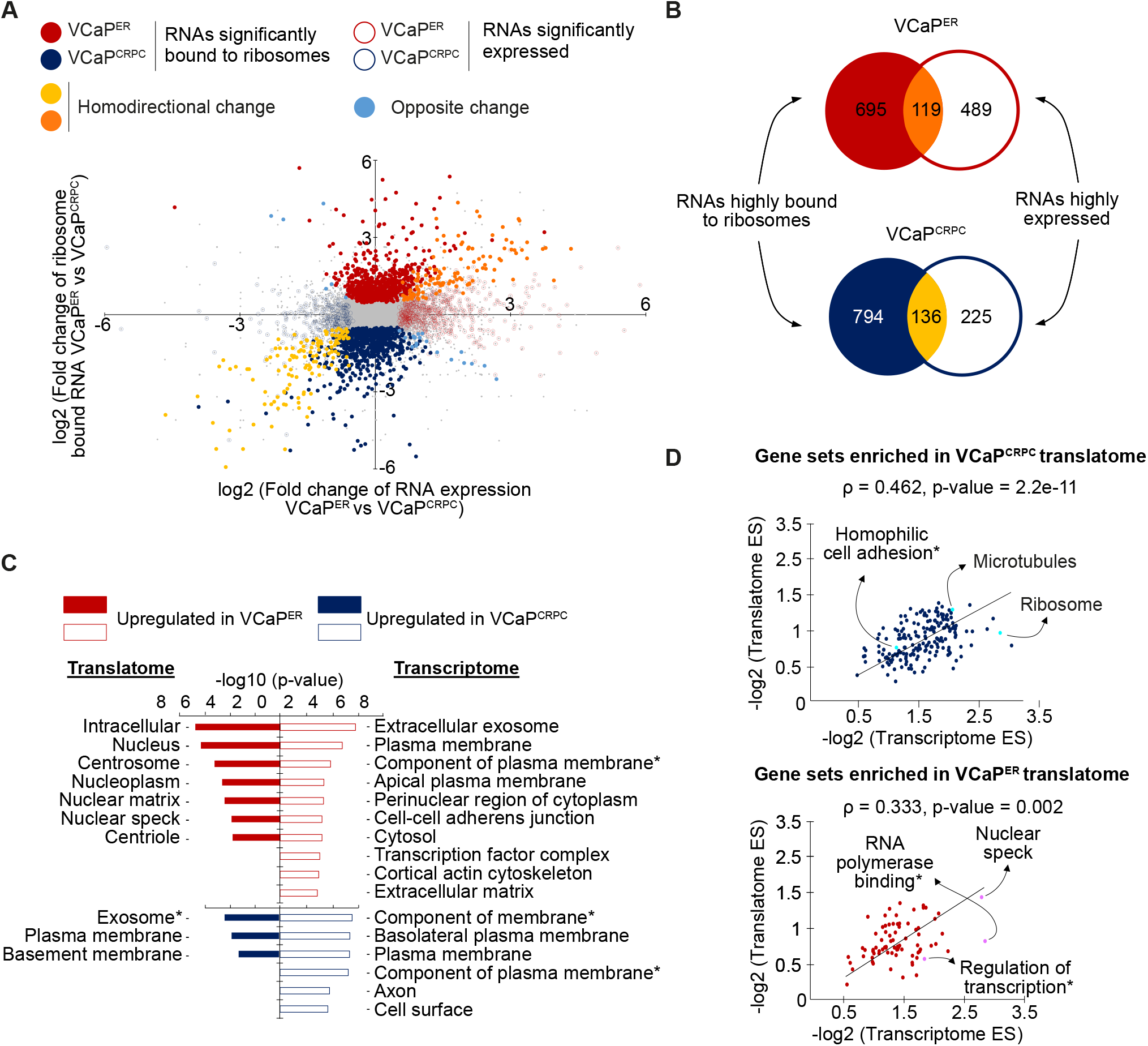
Transcriptome and translatome analysis highlights variations in translated genes in VCaP^ER^. **(A)** Scatterplot plot and **(B)** Venn diagram highlighting RNAs significantly up or downregulated (empty circles) or bound to ribosomes (full circles) in VCaP^ER^ (red) compared to VCaP^CRPC^ (blue). Significant genes are colored (adjusted p-value <0.05) (n=2 RNA-seq replicates for polysome profiling; n=2 RNA-seq replicates for total RNA sequencing). **(C)** GO term enrichment analysis shows top 10 significant cellular component terms (GOCC) enriched for genes upregulated in the translatome or transcriptome of VCaP^ER^ (red) or VCaP^CRPC^ (blue). **\*** Full names of GO terms: extracellular exosome, integral component of plasma membrane, integral component of membrane. **(D)** Scatterplots of GSEA enrichment scores (ES) for gene sets enriched in the VCaP^CRPC^ (top) and VCaP^ER^ (bottom) translatome. ES scores in VCaP^CRPC^ (blue) or VCaP^ER^ (red) translatomes and transcriptomes are plotted. Spearman correlation coefficients (ρ) and linear regressions (black lines) are indicated. *Full names of gene sets: RNA polymerase core enzyme binding, regulation of transcription elongation from RNA polymerase II promoter, homophilic cell adhesion via plasma membrane adhesion molecules.

To corroborate these results, we performed gene set enrichment analyses (GSEA) comparing the translatome with the transcriptome in either VCaP^ER^ or VCaP^CRPC^. We again observed cytoplasmic and membrane-linked gene sets enriched in total and polysomal RNA of VCaP^CRPC^, but also in total RNA for VCaP^ER^ (Supplemental Fig. S2B-D; Supplemental Table S4). In contrast, polysomal RNA in VCaP^ER^ was enriched for gene sets involved in nuclear processes such as transcription and RNA polymerase binding (Supplemental Fig. S2E; Supplemental Table S4). Furthermore, enrichment scores (ES) for gene sets enriched in the translatome of VCaP^ER^ show only a moderate correlation (ρ = 0.333) with ES for gene sets enriched in the transcriptome of VCaP^ER^ (Fig. 2D). However, in VCaP^CRPC^, ES for gene sets enriched in the translatome correlated relatively well with ES corresponding to the transcriptome (ρ = 0.462). Again using GSEA, but this time considering gene sets that were enriched in the transcriptomes of VCaP^CRPC^ and VCaP^ER^, we found a strong correlation between ES in the translatome and transcriptome for both cell lines (ρ = 0.457 in VCaP^CRPC^, ρ = 0.489 in VCaP^ER^) (Supplemental Fig. S2F). This suggests that gene sets or pathways enriched within the translatome of the ENZ-resistant cell line VCaP^ER^ are unique to these cells and differ from both their transcriptome, and the translatome and transcriptome of VCaP^CRPC^. Taken together, these results are indicative of perturbations in the translational landscape during ENZ-resistance acquisition. This could potentially cause changes in the expression of specific proteins and may therefore participate in promoting the resistant phenotype exhibited by VCaP^ER^ cells.

### ENZ-resistant cells exhibit deregulated protein expression of translation regulators

To understand how changes in ribosome association affect the resulting protein levels in ENZ-resistant cells, we analyzed differential expression of total proteins. Using MS, we identified 2548 proteins of which 485 were differentially expressed between ENZ-sensitive and resistant cell lines (Fig. 3A; Supplemental Fig. S3A-C; Supplemental Table S2). To further validate that the proteins highly expressed in VCaP^ER^ were linked to ENZ-resistance globally and not specific to our model, we quantified the proteome of another ENZ-resistant cell line (MR49F), which was derived from LNCaP cells (Bishop et al. 2017). We show that proteins highly expressed in VCaP^ER^ corresponded well with those highly expressed in MR49F, which was not the case for downregulated proteins (i.e. highly expressed in VCaP^CRPC^) (Supplemental Fig. S3D, E). We therefore focused on this set of differentially expressed proteins as a starting point to identify gene pathways or biological processes implicated in ENZ-resistant PCa. Hence, we performed a gene network analysis and found that ENZ-resistance promoted up-regulation of two main clusters: mitochondrial translation factors, with multiple mitochondrial ribosomal proteins upregulated, as well as mitochondrial electron transport, with numerous subunits of the NADH: ubiquinone oxidoreductase complex (mitochondrial respiratory complex I) (Fig. 3B; Supplemental Fig. S4A; Supplemental Table S5). Additionally, four main clusters were observed for proteins downregulated in VCaP^ER^, showing involvement in DNA-dependent DNA replication, translational initiation, mRNA splicing and post-translational protein modification (Fig. 3C; Supplemental Fig. S4B; Supplemental Table S5). Most notably, various core translation regulators such as EIF4B and EIF4EBP1, and ribosomal proteins such as RPL9, RPL11, RPS13 and RPS24, which have already been linked to malignant PCa (Mangangcha et al. 2019; Arthurs et al. 2017; Hernández et al. 2019; Hsieh et al. 2015), were found to be downregulated in VCaP^ER^ (Fig. 3C). These results were corroborated by GSEA (Supplemental Fig. S4C; Supplemental Table S6) and together, suggest that ENZ-resistance affects overall translation and promotes a switch from cytoplasmic to mitochondrial translation.

**Figure 3.**
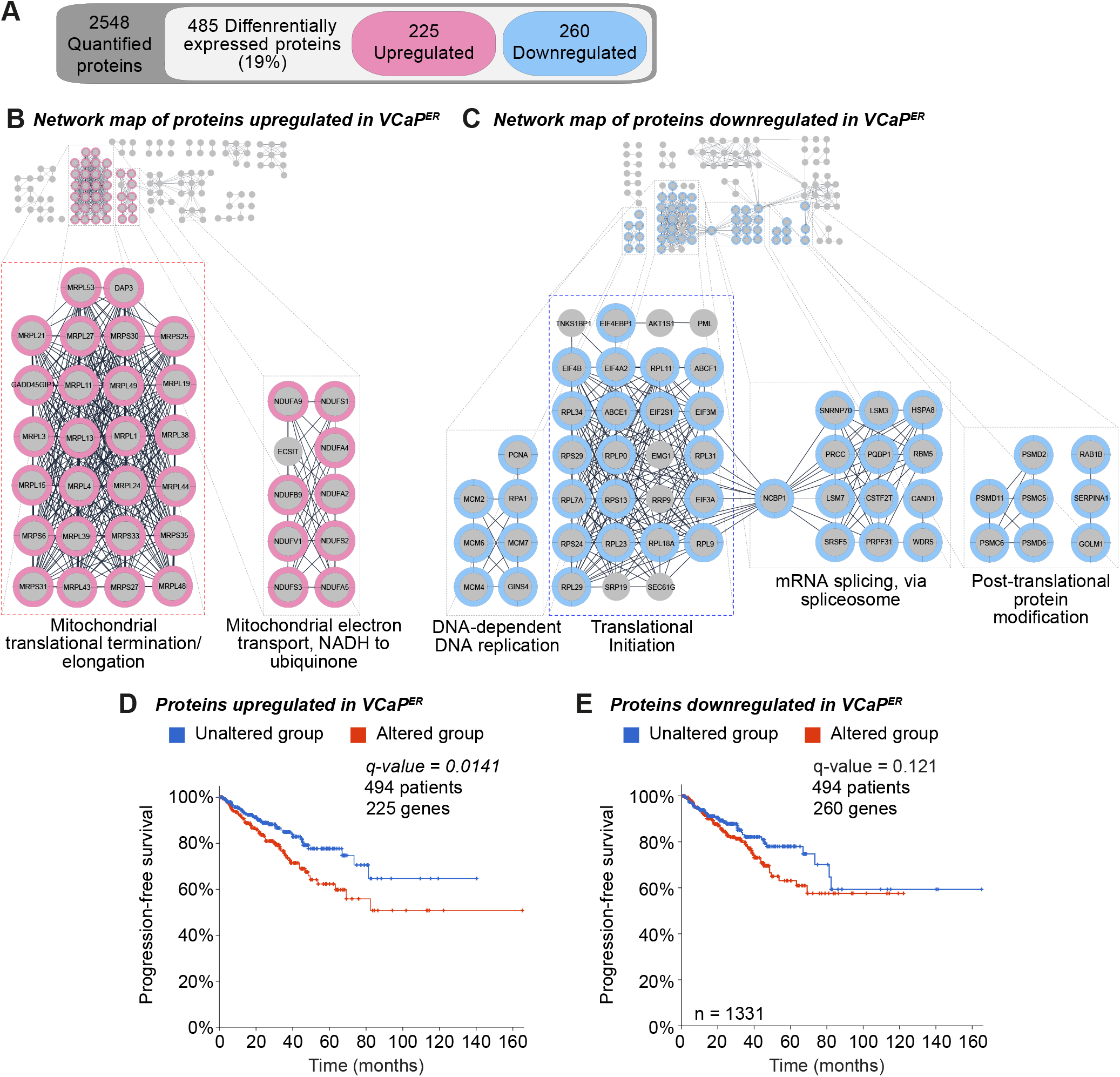
Changes occur in the proteomic landscape upon resistance acquisition. **(A)** Mass spectrometry results show differentially expressed proteins in VCaP^ER^ cells compared to VCaP^CRPC^ **(B)** Network analysis shows clusters formed by proteins up or **(C)** downregulated in VCaP^ER^. Main clusters are identified and highlighted in pink (upregulated clusters) and blue (downregulated clusters). A switch between cytoplasmic and mitochondrial translation-associated networks is highlighted by dashed frames (blue for cytoplasmic and red for mitochondrial translation). **(E)** Kaplan-Meier graph of progression-free survival for patients according to alterations in genes coding for proteins up or **(F)** downregulated in VCaP^ER^.

To evaluate the correlation between proteins enriched in our ENZ-resistance model and patient tumor data, we explored publicly available data sets through cBioPortal (Cerami et al. 2012; Gao et al. 2013). However, due to lack of available ENZ-resistance proteomic datasets and the fact that PCa is a cancer driven by copy number alterations (Fraser et al. 2017), we opted to correlate our data to genomic alterations identified in PCa patients, which consisted mainly of gene amplifications or deep deletions (Supplemental Fig. S5A). As such, we investigated copy number variations occurring in genomic loci for differentially expressed proteins from our model, in relation to patient clinical outcomes in the TCGA dataset. We found that these genomic alterations for proteins upregulated in VCaP^ER^ corresponded to significantly lower disease-free survival in PCa patients (Supplemental Fig. S5B). This difference between the altered and unaltered groups was also apparent with downregulated proteins but to a much lesser extent (Supplemental Fig. S5C). However, a stronger link can be established between worse progression-free survival and copy number variations in the VCaP^ER^ upregulated proteins, but not with the downregulated proteins (Fig. 3D, E). Interestingly, for all differentially expressed proteins, either up- or downregulated in VCaP^ER^, significant correlation was found with low overall patient survival (Supplemental Fig. S5D, E) and high Gleason score (Supplemental Fig. S5F, G). This suggests that focusing on the genomic alterations of VCaP^ER^ upregulated proteins can distinguish between overall and disease/progression-free survival of patients. We also attempted to validate our data to address the effects of alterations on gene expression; however, analysis of genomic and transcriptomic data from TCGA samples revealed that copy number variations were not always accompanied by changes in RNA expression (Supplemental Fig. S6A, B). This implies that the association of the identified genes with PCa severity and/or drug resistance could indeed be independent of transcription and RNA levels, and hence depend on downstream processes such as post-transcriptional and translation regulation. Altogether, these patient data suggest that all differentially expressed proteins from VCaP^ER^ and VCaP^CRPC^ may be implicated in overall cancer aggressiveness. However, only the proteins upregulated in VCaP^ER^ correlate with progression-free survival, which may indicate a link with the development of cancer resistance, causing recurrence in PCa patients.

### Enzalutamide resistance coincides with decreased translation efficiency

To further explore the origins of the altered protein landscape observed in ENZ-resistant cells, we focused our analysis on translation efficiency (TE) of RNAs, which is determined by dividing read counts per million mapped reads (CPM) of RNAs in polysome profiling by the CPM of the corresponding genes in total RNA-seq (CPM_Polysome profiling_/CPM_Total RNA_) in VCaP^ER^ and VCaP^CRPC^ cells. We then calculated the TE ratio for each gene by taking the fold change in TE values between VCaP^ER^ and VCaP^CRPC^ (Supplemental Table S7) to evaluate the global effect of ENZ-resistance on translation. We found a negative shift in TE ratios in genes with significant differential TE values (DTE) between VCaP^ER^ and VCaP^CRPC^ cells (Fig. 4A; median value of −0.39) suggesting that ENZ-resistance has a negative impact on overall translation. Moreover, GO term analysis of genes exhibiting a high TE ratio shows an enrichment in nuclear GO terms (e.g.: nucleus, nucleoplasm, regulation of transcription, DNA replication and repair), whereas low TE ratio genes show enrichment for membrane or cytoplasmic GO terms (e.g.: membrane, cytosol, cell-cell adhesion, endosomal transport) (Fig. 4B; Supplemental Fig. S7A, Supplemental Table S8). This reflects our previously explored GO terms in the VCaP^ER^ and VCaP^CRPC^ translatome and underscores an ENZ-resistance induced reduction in translation efficiency.

**Figure 4.**
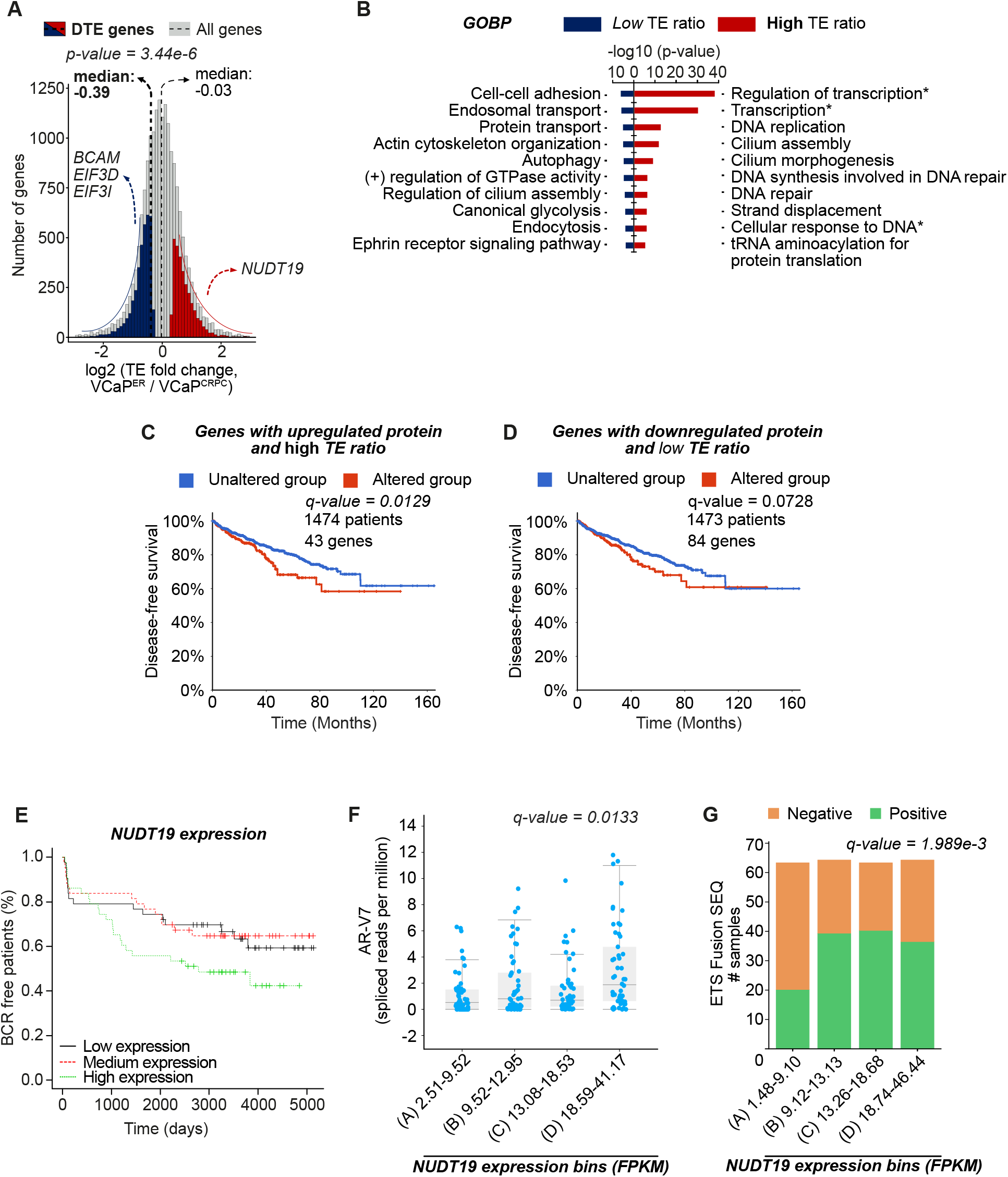
Enzalutamide resistance is accompanied by shifts in translation efficiency. **(A)** TE ratio distribution for all genes (pale grey) or genes with significant DTE in VCaP^ER^ compared to VCaP^CRPC^ (red for higher and blue for lower TE ratio). Medians are marked with dashed lines (thin lines for all genes, bold line for DTE genes). **(B)** GO term enrichment analysis shows top 10 significant biological process terms (GOBP) enriched in genes with low (blue) or high (red) TE ratios in VCaP^ER^ compared to VCaP^CRPC^. *Full names of GO terms: regulation of transcription, DNA-templated, transcription, DNA-templated, cellular response to DNA damage stimulus, **(C)** Kaplan-Meier graph of disease-free survival for PCa patients according to alterations in genes with either high or **(D)** low TE ratios. **(E)** Percentage of biochemical recurrence (BCR) for PCa patients expressing high, medium, or low protein levels of *NUDT19* in tissue microarrays. (F) Boxplot of *AR-V7* expression according to *NUTD19* expression in PCa patients. **(G)** Occurrence of ETS fusion according to *NUDT19* expression in PCa patients.

To identify potential biomarkers of PCa ENZ-resistance, we combined TE ratio results with our proteome data and focused on genes that possessed both differential protein expression as well as a corresponding differential TE ratio (Supplemental Fig. S7B). We found several candidate genes with high TE ratio (i.e higher TE in VCaP^ER^), which are implicated in metabolic processes, such as *NUDT19* (Nudix hydrolase 19) or *SHMT1* (Serine Hydroxymethyltransferase 1), whereas genes with lower TE ratios were implicated in cellular adherence, for example: *BCAM*, or translation initiation, such as *EIF3D* and *EIF3I* (subunits of the eIF3 core translation initiation factor). To evaluate their association with PCa clinical features, we again used TCGA patient genomic alteration data to correlate to patient related outcome. We observed that alterations such as copy number variations in genes from both categories (either up or downregulated TE ratios and protein expression in VCaP^ER^) were associated with significantly lower overall patient survival and were frequent in intermediate to high grade PCa (Supplemental Fig. S8A-D), underlining their potential role in PCa severity. Lower disease-free survival, however, was observed exclusively in the case of alterations in genes with high TE ratios and high protein expression in VCaP^ER^ (Fig. 4C, D). This highlights the fact that while genes with high protein expression and TE in either VCaP^ER^ or VCaP^CRPC^ are generally linked to higher cancer aggressiveness and lower patient survival, only those associated with VCaP^ER^ seem to be connected to disease recurrence in patients. Moreover, only some genes show significant changes in RNA expression according to copy number variation (Supplemental Fig. S9A, B). For example, *SHMT1* and *NUDT19* exhibit high TE ratios in VCaP^ER^, but show no significant changes in mRNA expression in cases of copy number variations. This highlights the fact that post-transcriptional mechanisms such as altered translation could potentially explain the link between these genes and decreased patient survival.

In order to see if the identified candidate genes with high TE ratios could act as potential biomarkers, we investigated NUDT19, a protein involved in RNA de-capping (Song et al. 2013) which has not yet been investigated in PCa and whose expression was validated in the ENZ-resistant MR49F cell line (Supplemental Fig. S10A). Using patient tissue microarray analyses, we show that high NUDT19 expression corresponds with earlier biochemical recurrence (BCR) post-prostatectomy (Fig. 4E). Furthermore, *NUDT19* expression correlated with a higher expression of the androgen receptor splice variant *AR-V7* (Fig. 4F), a high AR score in patients (Supplemental Fig. S10B) and a higher occurrence of the PCa-specific ETS fusion (Tandefelt et al. 2014) (Fig. 4G). Changes in *NUDT19* expression are also observed to be linked to significant increases in both the total fraction of the genome that is altered and the total mutation count in PCa samples, as well as inversely correlated to neuroendocrine PCa (NEPC) markers (Supplemental Fig. S10C-F). Together, these results reveal that incorporating proteome, translatome and transcriptome data can lead to the discovery of potent novel biomarkers for PCa severity or resistance.

### ENZ-resistance is linked to aberrant long non-coding RNA association to ribosomes

Following our investigations demonstrating the perturbation of protein-coding genes in ENZ-resistance, we next sought to investigate noncanonical associations with ribosomes. Indeed, many groups have put forward that cancer cells translate peptides noncanonically, promoting tumor initiation or growth (Wu et al. 2020; Sriram et al. 2018; Sendoel et al. 2017; Schuster and Hsieh 2019). We therefore first looked at ribosome association for non-coding genes in our ENZ-resistance model. We found that, while coding genes show no difference in association to ribosomes compared to what is expected, lncRNAs and processed transcripts were significantly over-represented in the VCaP^ER^ heavy polysome bound RNA (Supplemental Fig. S11A). We investigated if the previously observed global decrease in TE (Fig. 4A) occurred in both coding and non-coding genes. Interestingly, while a similar decrease could be observed for mRNAs (median = −0.41 for mRNAs with DTE), lncRNAs on the other hand, showed an inverse pattern, with a generally higher TE ratio in VCaP^ER^ (median = 0.51 for lncRNAs with DTE) (Fig. 5A). Consequently, the number of lncRNAs with significantly higher TE ratios in VCaP^ER^ was higher than expected (Supplemental Fig. S11B). Taken together, these findings suggest that some lncRNAs may actually code for peptides or undergo aberrant translation in resistant PCa. To investigate this hypothesis, we searched for putative peptides produced from lncRNAs with higher TE ratios in VCaP^ER^ in our MS data of VCaP^CRPC^ and VCaP^ER^, as well as in other publicly available datasets (Chen et al. 2020a; Bazzini et al. 2014; Slavoff et al. 2013), resulting in 189 lncRNA-encoded peptides being detected (Supplemental Table S9). Of these lncRNA-encoded peptides, 41 exhibited higher TE ratios in VCaP^ER^, while 23 lncRNAs with low TE ratio were also detected (Fig. 5B). To evaluate the importance of these lncRNAs in PCa patient overall survival, we again used the TCGA PCa dataset, which contained data for 22 high TE lncRNAs and 12 low TE lncRNAs. We observed that high TE lncRNAs were subject to genomic alterations such as copy number variations much more frequently than low TE lncRNAs in prostate tumors (Supplemental Fig. S12A). By investigating patient survival in cases of alterations for these lncRNAs, we found that gene amplifications or deep deletions for both lncRNAs with high and low TE ratios could correspond to poor overall patient survival (Supplemental Fig. S12B). In fact, of the 22 lncRNAs with high TE ratios, 11 showed a strong negative correlation with overall survival (Fig. 5C, D; Supplemental Fig. S12C), whereas alterations in only 3 out of the 12 lncRNAs with low TE ratios correlated with poor patient outcome (Fig. 5D; Supplemental Fig. S12D, E). This is a significantly lower fraction compared to high TE ratio lncRNAs suggesting that lncRNAs with high TE detected in VCaP^ER^ are more likely to be copy number altered in resistant PCa. We then investigated if alterations for these lncRNAs could also be linked to PCa grade. Analysis of Gleason scores from PCa patients revealed a high prevalence of alterations in intermediate to high grade cancer, most notably in lncRNAs with high TE ratios but to a lesser extent for low TE ratio lncRNAs (Fig. 5E; Supplemental Fig. S12F). Interestingly, for some of the identified lncRNAs, expression of the transcript was significantly altered upon genomic alterations, (Supplemental Fig. S13A, B). For example, the expression of lncRNAs such as *CRNDE*, *OIP5-AS1* (also known as *Cyrano*) and *JPX* vary following the type of alteration. Other lncRNAs such as *LINC00467* and *TMEM147-AS1* do not seem to be affected, in terms of RNA expression, by copy number variations, and a higher translation efficiency for these lncRNAs might therefore explain their link with PCa and drug resistance. These findings underscore a novel link between lncRNA association to ribosomes and high grade or resistant PCa.

**Figure 5.**
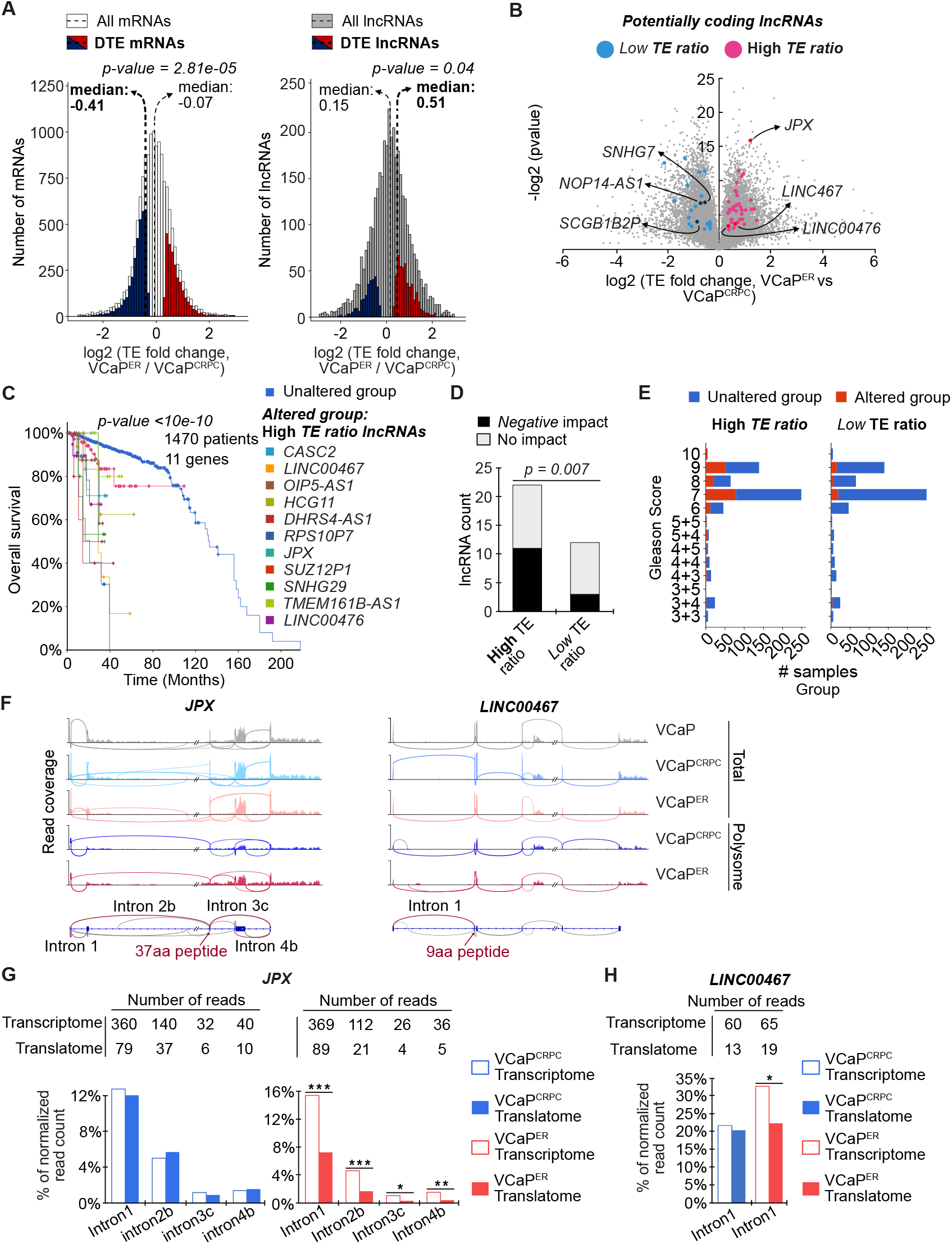
Aberrant association of lncRNAs to ribosomes in PCa drug resistance. **(A)** TE ratio distributions for all enes, or for genes with differential translation efficiency (DTE) in VCaP^ER^ compared to VCaP^CRPC^ (red for higher nd blue for lower TE ratio) for mRNAs and lncRNAs. Medians are marked with dashed lines (thin lines for all enes, bold line for DTE genes). **(B)** Volcano plot of lncRNAs with high (pink) and low (blue) translation fficiencies in VCaP^ER^, which were also detected in MS datasets. **(C)** Kaplan-Meier graph showing overall survival for PCa patients according to alteration of single lncRNAs with high TE ratios, correlating to poor atient outcome. **(D)** Quantification of lncRNAs with either high or low TE ratios, whose alteration positively or negatively correlate with PCa patient survival. **(E)** Distribution of Gleason scores for patients according to lterations in lncRNAs with high or low TE ratios in VCaP^ER^. **(F)** Sashimi plots for two candidate lncRNAs in PCa cell lines showing read coverage and splice junction usage. Schematics show potential alternatively spliced introns (red) and putative peptides for *JPX* (left) and *LINC00467* (right). **(G)** Quantification of *JPX* or **(H)***LINC00467* lncRNA split reads count f or selected splicing events, corresponding to alternatively spliced transcripts in the transcriptome and translatome of VCaP^CRPC^ (blue) or VCaP^ER^ (red) (*: p < 0.05, **: p < 0.01, **: p < 0.001).

We next asked what drives a nuclear lncRNA, such as *JPX*, to be shuttled into the cytoplasm to be translated. One possibility resides in the production of alternative isoforms for these lncRNAs, for which the subcellular localization could be cytoplasmic, a process under increasing scrutiny for various RNAs (Zeng and Hamada 2020; Yoshimoto et al. 2017), which could lead to the production of peptides. We found that some lncRNAs with high TE ratios such as *JPX*, *LINC00467* and *WARS_AS1*, showed several events of alternative splicing in VCaP^ER^ (Fig. 5F-H; Supplemental Fig. S14A, B). Moreover, *JPX* presents several variations in exon choice between the VCaP^ER^ translatome and transcriptome (Fig. 5G; Supplemental Fig. S14B). Interestingly, these changes are absent from VCaP^CRPC^ and may therefore explain the increase in TE observed for *JPX* in VCaP^ER^. Another lncRNA, *LINC00467*, shows similar switches in isoform expression between VCaP^ER^ total and polysome-bound RNAs but not in VCaP^CRPC^ (Fig. 5H; Supplemental Fig. S14B). Interestingly, the putative peptide-coding sequences in *JPX* and *LINC00467* are on their fourth and second exons respectively and are both directly adjacent to splice junctions whose activities differ between VCaP^ER^ and VCaP^CRPC^ (Fig. 5F). Altogether, these results show that alternative splicing of lncRNAs could lead to splice variants which may differentially bind to ribosomes, to either be translated or affect translation of other genes. Existence of these lncRNA variants could explain the aberrant association to ribosomes which is observed in VCaP^ER^ and grant coding potential to otherwise non-coding genes in the context of PCa resistance.

## DISCUSSION

Genomic alterations in cancer are often viewed as the top of the hierarchy driving cancer biology through downstream transcription and subsequent translation to protein. However, it would be false to assume that changes on one of these three levels (genomic, transcriptomic, and proteomic) are necessarily direct and linear. Indeed, various regulatory mechanisms are responsible for finely controlling the processes that leads to mature proteins in our cells, and several of these regulatory steps are altered in cancer. Here, we report, to the best of our knowledge, the first detailed analysis of the translatome in drug resistant PCa, which we investigated using novel models of ENZ-resistance. We show that only 20% of RNAs significantly more transcribed in our drug resistant PCa model are also upregulated in the polysomal fractions (119 out of 608 genes). Additionally, both transcribed and polysome-associated RNAs only moderately correlate with proteins detected by MS, which suggests post-transcriptional regulatory mechanisms not detectable in genomic or transcriptomic data. Indeed, previous studies have also shown low correlation between the abundance of specific RNA transcripts and of their related protein in cancer (CPTAC et al. 2016; Zhang et al. 2016a, 2014). This is due, at least in part, to translational control, which remains one of the key checkpoints for regulating the expression of protein coding genes. Mounting evidence further shows an association between cancer resistance and perturbation of translation mechanisms, through upregulation or downregulation of certain translation factors and signalling pathways such as mTOR, *EIF4B* or eIF2 (Hernández et al. 2019; Murugan 2019). Recent publications have shown that affecting these pathways and regulators of translation is a promising target for cancer therapy (Hua et al. 2019; Hernández et al. 2019). We found that the main pathways deregulated in ENZ-resistant PCa cells were indeed implicated in translation, namely translation initiation and mitochondrial translation. We show that the main genes upregulated in the context of PCa drug-resistance are genes affecting mitochondrial translation and metabolism, for example mitochondrial ribosomal proteins and subunits from the mitochondrial respiratory complex I. This is consistent with previous data reporting clinical therapeutics targeting mitochondrial processes such as Gossypol, the G3139 anti-sense oligonucleotide and 2-deoxy-d-glucose (Hsu et al. 2016; Yang et al. 2016; Kim et al. 2017; D’Souza and Minczuk 2018; Fulda et al. 2010; Zhang et al. 2011). Our model highlights the dysregulation of certain ribosomal genes, for example *RPL9* and *RPS13*, and translation regulators such as *EIF4B* and *EIF4EBP1*, which have already been found to be implicated in PCa through the use of varied models (Hsieh et al. 2015; Verma et al. 2019; Liu et al. 2019; Guo et al. 2011; Ren et al. 2013). Furthermore, the downregulation of genes implicated in translation may suggest that development of drug resistance in PCa could benefit from this reprogramming of translation. Future work will need to explore if and how current therapeutics targeting translation could affect the emergence of drug resistance in castration resistant PCa. Indeed, all of the widely used therapies currently employed to treat castration resistant prostate cancer ultimately lead to drug resistance (Buttigliero et al. 2015), and novel avenues of research therefore need to be explored. Studying the differences between castration resistance and drug resistance could therefore guide the discovery of therapeutic targets that would overcome the otherwise inevitable development of the drug resistant state of this disease.

With an abundance of information from transcriptomics and genomics PCa data now available, the relative paucity of detailed and specific data available on specific translational changes is remarkable. Previous studies have demonstrated that focusing only on either genomic or transcriptomic data for instance, paints an incomplete picture of cancer, and that work to integrate multiple types of data is necessary for a better understanding of the disease (Sinha et al. 2019). Proteome and translatome data are hence essential for integrative approaches to discover novel biomarkers and therapeutic targets. We demonstrate a pipeline for discovery of such potential new targets for drug resistant PCa, through the combined transcriptomic, translatomic and proteomic data from resistance models and publicly available genomic data. We discovered several genes with altered translation efficiencies in the context of resistance. Interestingly, we found that in our model, genes with a reduced association to ribosomes and protein abundance in resistant cells still correlate with a higher cancer grade and worse overall survival. This is most likely related to the fact that we compare drug resistant cells to castration resistant PCa cells. Tumor castration resistance in patients already corresponds to a severe stage of the disease, and genes upregulated in this stage are understandably already linked to poorer patient prognosis. The novelty in our approach resides in the identification of a signature specific to ENZ-resistance, which is composed of genes linked to cancer recurrence after treatment, as shown by our analyses of disease-free survival and progression-free survival. These genes may be used in the future as markers to guide therapeutic options. Out of the identified target genes, *NUDT19* represents a highly interesting candidate, with elevated protein levels in resistant PCa and whose expression or alteration is linked to various determinants of high grade and resistant PCa. *NUDT19* is part of a gene family involved, among other things, in mRNA decapping, a process highly relevant for translation regulation as it renders mRNAs repressed or degraded (Song et al. 2013). Further research will need to assess if affecting *NUDT19* expression could represent a therapeutic target for combatting enzalutamide resistance, but these early results are encouraging as to its potential value as a biomarker for PCa drug resistance.

Interestingly, our study also shows that translation perturbation is not limited to protein coding genes in drug resistant PCa, but also affects non-coding genes. Indeed, while it is well known that disruptions in the regulation of coding genes are important in cancers, recent evidence has shown that non-coding genes can also play a major role (Schmitt and Chang 2016). In fact, we found that translation efficiency is generally negatively affected in protein coding genes, which may be explained by the downregulation of translation factors (such as EIF4B or EIF4EBP1) in VCaP^ER^ cells; but this does not explain the overall increase in ribosome-bound lncRNAs and the positive shift in translation efficiency observed for these otherwise non-coding transcripts. It has been shown that lncRNAs may bind ribosomes, and some have been shown to produce functional peptides (Wu et al. 2020; Chen et al. 2020a; Ruiz-Orera et al. 2014). We show that for some lncRNAs, different splice isoforms are bound by ribosomes in resistant cells when comparing to sensitive cells, which is consistent with the previously observed deregulation of splicing (Fig. 3C). This suggests that differences in the choice of lncRNA splice isoforms in drug-resistant cells could affect the function of these lncRNAs. Furthermore, a recent study found multiple of our identified lncRNAs (such as *JPX*, *LINC00467* and *CASC2*) bound by PCa-linked alternative splicing regulators in LNCaP cells, which could play a role in the alternative splicing observed in VCAP^ER^ (Fei et al. 2017).

Our study also highlights that subcellular compartmentalization of lncRNAs could play a role in PCa drug resistance. For example, *JPX*, a well-known nuclear lncRNA implicated in X-chromosome inactivation (Tian et al. 2010; Sun et al. 2013) was found in the ribosomal RNA fractions in our polysome profiling data, as well as in other datasets (Chen et al. 2020a; Bazzini et al. 2014; Slavoff et al. 2013). This association to ribosomes indicates that *JPX* transcripts can exit the nucleus, which contrasts what has previously been reported. One possibility that would reconcile the mostly nuclear localization of *JPX* and other nuclear lncRNAs with ribosome binding lies upon the alternative splicing of RNAs, which often results in multiple isoforms with possibly diverse cellular localizations (Yoshimoto et al. 2017; Zeng and Hamada 2020). Indeed, for some lncRNAs highlighted here, significant association to ribosomes of specific splice variants was found exclusively in VCaP^ER^, hinting at potential mechanisms of nuclear export or ribosome binding exclusive to these isoforms. Whether the observed alterations of RNA splicing may also be linked to higher expression of the AR-V7 splice variant is topic for further research. Moreover, while ribosome binding does not guarantee translation of an RNA, several peptides corresponding to *JPX* and the other identified lncRNAs are detected by MS in diverse cellular contexts. However, the question remains as to whether these lncRNAs produce stable and functional peptides in drug resistant PCa cells.

Nonetheless, we show that genomic alterations in select lncRNAs with potential coding capacity strongly correlate with low patient survival and high PCa grades, underscoring their importance as potential biomarkers and therapeutic targets. In some cases, copy number variations for these lncRNAs has been linked to changes in RNA expression in patient samples suggesting that expression as well as alterations of these lncRNAs could serve as novel prognostic biomarkers for prostate cancer drug resistance. We therefore, for the first time, link the ribosome binding and possibly the coding potential of these lncRNAs to PCa drug resistance. Our data corroborates previous studies linking some of our candidate lncRNAs to PCa (Gao et al. 2019; Zhang et al. 2016b; Wan et al. 2015) and highlighting potential novel roles for several others. Using lncRNAs for drug resistant PCa early detection and treatment represents an avenue of high interest due to their highly restricted spatio-temporal expression patterns and their relative ease of targeting, for example using specific anti-sense oligonucleotides (Nandwani et al. 2020; Chen et al. 2020b; Bonetti and Carninci 2017). While research on cancer therapies targeting lncRNAs has not yet reached clinical trials, several studies have shown promise and widening the knowledge on lncRNAs of potential clinical importance therefore constitutes a priority for the coming years. This new information could in turn lead to better targeted therapies and to a greater ease of detection for drug resistant PCa.

In conclusion, our study highlights the occurrence of translation dysregulation during the development of PCa drug resistance. We reveal an unusual shift in ribosome binding from protein coding genes to lncRNAs in ENZ-resistant PCa cells. However, several questions remain to be answered: Is this remodeling a consequence or driver of drug resistance? Do these lncRNAs code for functional peptides or regulate translation, or can the fact that they bind to ribosomes lead to PCa drug resistance? Our study brings forward novel concepts and prognostic biomarkers that relate to the translation output of drug resistant cancer cells and enables the discovery of potential biomarkers hidden from previous transcriptome and proteome analyses.

## METHODS

### Cell culture and drugs

LNCaP and VCaP cell lines were obtained from ATCC. All cell lines were cultivated at 37°C with 5% CO_2_. LNCaP and MR49F (ENZ resistant derivated from LNCaP) were cultivated in RPMI 10% FBS whereas VCaP and VCaP^ER^ (ENZ resistant derivated from VCaP) were cultivated in DMEM 10% FBS with 1mM sodium pyruvate. ENZ-resistant cell lines were maintained in 10 μM ENZ. ENZ (MDV3100) purchased from MedChemExpress (Cat. No.: HY-70002).

### Generation of enzalutamide-resistant cell lines in mice

All animal procedures were performed according to Canadian Council on Animal Care guidelines and with approval of the Animal Care Committee of the University of British Columbia (protocol # A12-0210). One million VCaP cells were inoculated on both flanks of six-week-old male athymic nude mice (Harlan Sprague-Dawley, Inc). Two weeks later, when tumors reached an approximate volume of 200mm^3^, mice were surgically castrated. Castration resistance subsequently developed and when these tumors were growing beyond their pre-castration size, tumors were freshly harvested, washed, passaged and isolated from stromal cells in RPMI with 10% fetal bovine serum. Among several tumors concurrently passaged this way, cells termed VCaP^CRPC^ were selected for further experiments. Mice with castration-resistant tumor were then force-fed with 10mg/kg ENZ (or vehicle) 5 days per week until tumor recurrence, at which point cells termed VCaP^ER^ were isolated as previously described, and maintained in medium supplemented with 10μM ENZ. See Figure 1A.

### Statistical analysis

Statistical significance of differences among groups was determined by two-tailed unpaired and paired Student’s t-test as well as analysis of variance (ANOVA) with Student-Newman-Keuls post hoc analysis or Kruskal-Wallis test with Dunn’s multiple comparison test when assumption for equal variances could not be met, using Sigma Stat (SPSS) or PRISM 5.0 (GraphPad). Correlation analyses employed Pearson correlation coefficients, or Spearman correlation coefficients when assumption of equal variances could not be satisfied. Analysis of expected and observed frequencies was accomplished via Chi-squared tests. Differences with P < 0.05 were considered statistically significant.

### Proliferation of ENZ resistance cell model

Cell proliferation was measured using Cell-Counting Kit-8 (CCK8) (Dojindo, Gaithersburg, MD) which quantifies cellular dehydrogenase activity. Briefly, cells were seeded into 96-well plates and treated with 0.5 to 50 μM of ENZ. After 7 days, CCK-8 solution was added in media for a final concentration of 10% and incubated at 37°C for 3 hours. Cell growth was determined by optical density (OD) measurements at 450 nm with TECAN Infinite F50. Calculations were performed according to manufacturer’s instructions. Experiment was repeated 3 times with each cell line.

### Western blot for AR and AR-V7

VCaP^CRPC^ and VCaP^ER^ cells were lysed in RIPA buffer containing protease and phosphatase inhibitors. Total proteins were separated by SDS-PAGE and transferred on nitrocellulose membrane. After blocking with 5% milk, membranes were blotted overnight with primary antibodies against AR, AR-V7 or ACTB as loading control (Cell signaling technologies) diluted 1:1000. Membranes were then incubated with HRP-conjugated secondary antibodies (Jackson ImmunoResearch Laboratories Inc, West Grove, PA) 2 hours in 5% milk (1:10 000). Proteins were imaged on Chemidoc MP (Bio-Rad, Hercules, CA) with ECL reagent.

### Polysomal profiles and isolation of polysome-associated RNAs

VCaP^CRPC^, VCaP^ER^, LNCaP and MR49F cells were grown as described above, in 100-mm tissue culture dishes to ~ 80% confluence. Only VCaP^ER^ and MR49F were treated with ENZ at 10 μM. Cells were scraped in 1 mL of polysomal buffer (20 mM Tris, pH 7.5, 150 mM NaCl, 1.25 mM MgCl_2_, 5 U/mL RNasin, cOmplete™ EDTA-free Protease Inhibitor Cocktail (Roche, Indianapolis, IN), and 1 mM dithiothreitol), and Nonidet P-40 was added to a final concentration of 1% for lysis, 15 minutes on ice. Extracts were clarified by centrifugation at 12,000 g for 20 minutes at 4°C. RNA concentration was measured by spectrophotometry and ~20 OD_260_ units of RNA were loaded onto a 15-55% sucrose gradient. The gradients were centrifuged for 2.5 hours at 37,000 rpm (223 000g) (SW 40 TI Beckman rotor) and then placed on an Automated Density Fractionation System to collect fractions. Each fraction was collected into individual tubes with continuous monitoring of absorbance at 254 nm. Absorbance was recorded on chart paper to generate polysomal charts. RNA from each fraction was extracted by phenol-chloroform extraction and fractions corresponding to light or heavy polysomes were respectively pooled together.

### RNA extraction and library preparation for RNA sequencing

VCaP, VCaP^CRPC^, VCaP^ER^, LNCaP and MR49F cells were grown in 100-mm tissue culture dishes to ~ 80% confluence. Total RNA was extracted with Trizol reagent (Life Technology) and heavy polysomal RNA was prepared by phenol-chloroform extraction. RNA quality was verified with the TapeStation 4200 (Agilent Technologies). RNA libraries were made from 0.2 ug of RNA in accordance with the TruSeq stranded mRNA kit protocol (Illumina; # 20020594) and TruSeq RNA Single Indexes Set A and B (Illumina, # 20020492 and 20020493). Library qualities and sizes were checked with the TapeStation 4200 and then quantified using the KAPA Library Quantification Kit for Illumina platforms (Kapa Biosystems). Libraries were sequenced on the Illumina NextSeq500 sequencer to a depth of about 50 millions of 75-bp pair reads per library.

### Transcriptome and translatome RNA sequencing analysis

The reads were aligned to the GRCh37 human genome (Ensembl release 75) by STAR (v2.7.5) (Dobin et al. 2013). Read alignments were merged and disambiguated, and a single BAM (Binary Alignment Mapped) file output per library or sample was used. BAM files were then additionally filtered to remove reads with a mapping quality (MAPQ) less than 13, and all ribosomal and mitochondrial RNA reads. Alignments were assembled using Cufflinks (v2.2.1) (Trapnell et al. 2010) using the –g parameter to construct a genome annotation file against the reference gene model (Ensembl release 75) and to identify novel transcripts. Raw read counts were obtained by mapping reads at the gene level using the Cufflinks assembled transcript annotation file with the featureCounts tool from the SubRead package (version 2.0.0) (Liao et al. 2014) using the exon counting mode. EdgeR R-package (v3.12.1) (Robinson et al. 2010) was then used to normalize the data, calculate transcript abundance (as counts per million reads (CPM)) and perform statistical analysis. Briefly, a common biological coefficient of variation (BCV) and dispersion (variance) was estimated based on a negative binomial distribution model. This estimated dispersion value was incorporated into the final EdgeR analysis for differential gene expression, and the generalized linear model (GLM) likelihood ratio test was used for statistics, as described in EdgeR user guide. Genes were considered as significantly up or downregulated in either total or polysome-bound RNAseq if their fold change between VCaP^ER^ and VCaP^CRPC^ were superior to 1.25 fold, with a FDR ≤ 0.05. Analysis of TE values and significant differences between VCaP^ER^ and VCaP^CRPC^ were done by dividing CPM in the heavy polysome-bound RNA-seq by the corresponding CPM value in total RNA-seq. Differential TE ratio was calculated by dividing TE values from VCaP^ER^ by TE from VCaP^CRPC^, and significance was determined using R using the p.adjust function with the FDR settings. For analysis of RNA splicing, normalized counts for split reads corresponding to exon-intron junctions were quantified for select lncRNAs. Significance was determined by Chi-squared test comparing expected and observed frequencies in the translatome compared to the transcriptome. All statistical analyses and data visualization were done in R using R basic functions and the following packages: gplots (3.1.1), stats4 (3.5.1), plyr (1.8.4), dplyr (0.8.1), and ggplot2 (3.1.1). Raw RNA-seq data will be submitted to the NCBI Gene Expression Omnibus before manuscript acceptance, and is available upon request in the meantime.

### Sample preparation for Mass Spectrometry

Protein pellets were resuspended in 100 μL of 50 mM ammonium bicarbonate, 0.5% deoxycholate and sonicated on ice with a microprobe Sonic Dismembrator 550 (Fisher Scientific) as follow: 20 × 1 seconds at power 2 followed by 5 × 3 seconds at power 4. The extract was centrifuged at 20,817 g for 15 minutes at 4°C. The supernatants were transferred to new tubes and precipitated with acetone. The protein pellets were then resuspended in 100 μL of 500 mM triethylammonium bicarbonate, 0.5% deoxycholate. Protein concentrations of each sample was determined by colorimetric Bradford assay.

### Tryptic digestion and TMT labeling

10 μg of each sample was used for TMT labeling (Thermo Fisher Scientific). Proteins were denatured for 5 minutes at 95°C and then reduced with 50 mM TCEP for 30 minutes at 37°C before being alkylated with 100mM iodoacetamide for 30 minutes at room temperature in the dark. Samples were digested with 0.5 μg of trypsin (V5111; Promega) for ~15 hours at 37°C. After digestion, peptides were acidified to precipitate the deoxycholate and then purified with homemade C_18_ Stage-Tip before being lyophilized. The now dried peptides were dissolved in 30 μL of 100 mM triethylammonium bicarbonate and labeled with TMT 10-plex reagent (Thermo Fischer Scientific). Labeling was performed for 1 hour at room temperature and the reaction quenched with hydroxylamine for 15 minutes. The now labeled peptides were combined in one tube and speedvac to dryness without heat. Samples were cleaned up using solid-phase HLB cartridge (Water Corp.) before being speedvac to dryness.

### High pH - reverse phase fractionation

Peptides were fractionated into 14 fractions using a High-pH (pH 10) reversed-phase chromatography method using an Agilent 1200 HPLC system as previously described (Wang et al. 2011). The final fractions were dried and resuspended in 0.1% formic acid before mass spectrometry analysis.

### Mass spectrometry analysis

Approximately 1 μg of each fraction was injected and separated by online reversed-phase (RP) nanoscale capillary liquid chromatography (nanoLC) and analyzed by electrospray mass spectrometry (ESI MS/MS). The experiments were performed with a Dionex UltiMate 3000 nanoRSLC chromatography system (Thermo Fisher Scientific / Dionex Softron GmbH, Germering, Germany) coupled to an Orbitrap Fusion mass spectrometer (Thermo Fisher Scientific, San Jose, CA, USA) equipped with a nanoelectrospray ion source. Peptides were trapped at 20 μL/min in loading solvent (2% acetonitrile, 0.05% TFA) on a 5 mm × 300 μm C_18_ PepMap cartridge pre-column (Thermo Fisher Scientific / Dionex Softron GmbH, Germering, Germany) for 5 minutes. Then, the pre-column was switched online with a 75 μm × 50 cm Acclaim PepMap100 C_18_ - 3 μm column (Thermo Fischer Scientific/ Dionex Softron GmbH, Germering, Germany) and the peptides were eluted with a linear gradient from 5-40% solvent B (A: 0,1% formic acid, B: 80% acetonitrile, 0.1% formic acid) in 90 minutes at 300 nL/min. Mass spectra were acquired using a data dependent acquisition mode using Thermo XCalibur software version 3.0.63. Synchronous Precursor Selection-MS3 acquisition mode was used for this analysis. Full scan mass spectra (380 to 1500m/z) were acquired in the Orbitrap at a 60 000 resolution and using an AGC target of 2e5, a maximum injection time of 50 ms. Internal calibration, using lock mass on the m/z 445.12003 siloxane ion, was used. Precursors for MS2/MS3 analysis were selected using a TopSpeed of 3s. The most intense precursor ions were isolated in the quadrupole at 0.7 m/z, fragmented with 35% CID and the fragments detected in the ion trap. Following acquisition of each MS2 spectrum, an MS3 acquisition was performed by isolating of multiple MS2 fragment ions with a multi-notch isolation waveform (McAlister et al. 2014). MS3 analysis was detected in the Orbitrap at a 60 000 resolution after 45% HCD, with an AGC target of 1e5 and a maximum injection time of 120 ms. Dynamic exclusion of previously fragmented peptides was set for a period of 20 seconds and a tolerance of 10 ppm.

### Data analysis for mass spectrometry-based proteome quantification

Spectra acquired were processed using ProteomeDiscoverer 2.2 (Thermo). Files were searched against Uniprot (The UniProt Consortium 2017) homo sapiens protein database (93634 entries). Trypsin was set as enzyme and 2 missed cleavages were allowed. Deamidation (N, Q), oxidation (M), were set as dynamic modifications and carbamidomethylation (C), and TMT10-plex label (N-ter, K) were set as static modifications. Mass search tolerance were 10 ppm and 0.6 Da for MS and MS/MS respectively. For protein validation, a maximum False Discovery Rate of 1% at peptide and protein level was used based on a target/decoy search. MS3 spectra were used for quantification, with an integration tolerance of 10 ppm. Unique and razor peptides are considered for protein quantification and isotopic correction is applied on reporters. Data normalization was performed on total peptide amount. Peptides and protein result tabs were exported in Excel and means of three replicates per group were calculated. A fold change was calculated between the means of VCaP^ER^ and VCaP, VCaP^CRPC^ and VCaP, or VCap^ER^ and VCaP^CRPC^. Proteins or peptides with variations >>1.1 fold with a P ≤ 0.1 in either their VCaP^ER^/VCaP or VCaP^ER^/VCaP^CRPC^ fold changes were considered as significant (either up or downregulated), so long as no opposite variation was found in these two fold changes.

### Gene ontology and pathway enrichment analysis of differentially expressed genes

Gene Ontology (GO) term enrichment and Kyoto Encyclopedia of Genes and Genomes (KEGG) (Kanehisa et al. 2020; Kanehisa 2019; Kanehisa and Goto 2000) pathway enrichment were assessed using the Database for Annotation, Visualization and Integrated Discovery (DAVID 6.8) (Jiao et al. 2012). Biological processes (BP), cellular components (CC) and molecular functions (MF) annotations were analyzed. An FDR <0.1 and P < 0.05 were set as cut-off for significant enrichment.

### Construction of biological network

Interactions among identified differentially expressed proteins were mapped with the STRING database (Szklarczyk et al. 2018). Two protein-protein interaction (PPI) networks were constructed (for upregulated and downregulated proteins); experimentally validated interactions and databases with a required interaction score a 0.9. Subsequently, the PPI networks were imported into Cytoscape (Shannon et al. 2003) using stringApp (Doncheva et al. 2018). The identified hub genes related GO terms were used to construct a complete PPI network.

### Gene set enrichment, modules and network analysis

Gene Set Enrichment Analysis (GSEA) was performed with Broad Institute’s GSEA software (v4.1.0) (Subramanian et al. 2005). Expression data sets were created as text files according to GSEA specifications. We computed overlaps with the C2.cp.kegg (curated gene sets) and C5.go, C5.go.bp, C5.go.cc and C5.go.mf (GO gene sets) collections. Gene set permutations were performed 1000 times per analysis. An FDR <0.1 was set as cut-off for significant enrichment.

### Western blot for NUDT19

VCaP^CRPC^ and VCaP^ER^ cells were lysed in RIPA buffer containing protease and phosphatase inhibitors. Total proteins were separated by SDS-PAGE and transferred on PVDF membrane. After blocking with 5% milk, membranes were blotted overnight with primary antibodies against NUDT19 or TUBA1A as loading control diluted respectively in 1:10 000 and 1:10 000 (Abcam). Membranes were then incubated with HRP-conjugated secondary antibodies 2 hours in 5% milk (1:15 000). Proteins were imaged by revealing membranes on film with ECL reagent and quantified with the Image Studio Lite software.

### PCa patient cohort

This study was approved by the research ethics committee of CHU de Québec-Université Laval (Project 2016-2811). A sub-cohort of the CPC-GENE cohort (Fraser et al. 2017) was used for this study. The cohort comprised 136 men with intermediate risk PCa who underwent radical prostatectomy between 2005 and 2010 at CHU de Québec-Université Laval. Complete clinico-pathological data were available for all 136 subjects with a median follow-up of 11.8 years. Archived formalin-fixed and paraffin embedded specimens of radical prostatectomy from the 136 subjects were used to construct a tissue microarray (TMA).

### Immunohistochemistry on prostate tumor microarray (TMA)

Four micrometer-thick sections of the TMA blocks were first deparaffinized and rehydrated. Antigen unmasking was carried out in a DAKO PT-Link for 20 minutes at 97°C using EnVision FLEX buffer of appropriate pH for each antibody (Agilent). Then, peroxidases were inactivated in a 3% hydrogen peroxide solution. After incubation with a blocking solution, slides were stained overnight with antibody against Nudix Hydrolase 19 (NUDT19; 3.3μg/mL; Abcam) diluted in PBS containing 1% BSA. After washed, staining was revealed using the IDetect Super strain HRP polymer kit (Agilent) following manufacturer’s protocol. After DAB revelation, slides were counterstained with hematoxylin. Tissue samples were classified according to tumor staining intensity from 1 to 3. Intensity variation was also considered. Result for each patient is a mean of 3 cores. Results were confirmed by a second reader. Kaplan-Meier curves were established according to time to biochemical recurrence following prostatectomy.

## DATA ACCESS

All mass spectrometry files acquired in this study have been deposited to the MassIVE repository, assigned the MSV000086670 identifier and can be accessed at ftp://massive.ucsd.edu/MSV000086670/. The password to access them prior to publication is “prostate”. Processed data for all RNA sequencing experiments are available as supplemental tables. Raw RNA-seq data will be submitted to the NCBI Gene Expression Omnibus before manuscript acceptance, and is currently available upon request to the corresponding author.

## ACKNOWLEDGMENTS

We would like to thank Victoire Fort for comments and thorough editing of the manuscript. S.M.I. Hussein, P. Toren and J.-P. Lambert are Junior 1 Research Scholars of the Fonds de Recherche du Québec - Santé (FRQ-S). G. Khelifi is a recipient of training awards of the Fonds de Recherche du Québec - Santé (FRQ-S) and of the Natural Sciences and Engineering Research Council of Canada (NSERC). This work was supported by grants from the Canadian Institutes of Health Research (grants # PJT-378019 and PJT-168969) and the Cancer Research Society (23483), by an Early Investigator Award from the Canadian Urological Association Scholarship Foundation and by a Leader’s Opportunity Funds from the Canada Foundation for Innovation (36930, 37454, 41426).

## AUTHORS CONTRIBUTIONS

E.I.J.L. designed and performed experiments, analyzed and interpreted the data and wrote the manuscript, P.A. designed, optimized and performed the polysome profiling experiments, F.H.J designed and performed experiments, G.K. analyzed and interpreted the data and wrote the manuscript, V.S.G. designed, optimized and performed the polysome profiling experiments, A.Z. designed our cell-line derivation protocol and provided feedback on the manuscript, J.P.L. designed and analyzed M.S. experiments and wrote the manuscript, P.T. designed the ENZ-resistance model and other experiments and wrote the manuscript, R.M. designed polysome profiling experiments and wrote the manuscript, S.M.I.H designed experiments, analyzed and interpreted the data and wrote the manuscript.

