## Supplemental_material for "Prostate cancer resistance leads to a global deregulation of translation factors and unconventional translation of long non-coding RNAs"

### Table of Contents


| Table of Contents | Page |
|---|---|
| - **Supplemental Figure S1 :** Transcriptome and translatome analysis of ENZ-sensitive and resistant cell lines | 1 |
| - **Supplemental Figure S2:** Significant enrichment of gene sets in VCaPER and VCaPCRPC transcriptomes and translatomes | 3 |
| - **Supplemental Figure S3:** The VCaPER ENZ-resistance model recapitulates established features of drug-resistant PCa models. | 5 |
| - **Supplemental Figure S4:** Proteins up- and downregulated in VCaPER assemble in protein networks | 7 |
| - **Supplemental Figure S5:** Genomic alterations in VCaPER-specific proteins are linked to PCa drug resistance in patients | 9 |
| - **Supplemental Figure S6:** Genomic alterations are not always linked to RNA expression for genes coding for differentially expressed proteins in VCaPER | 11 |
| - **Supplemental Figure S7:** High TE ratio RNAs recapitulate key GO-terms from the translatome in VCaPER, but do not necessarily lead to higher protein expression. | 13 |
| - **Supplemental Figure S8:** Genomic alteration in proteins upregulated in VCaPER is linked to high PCa grade, independently of TE ratio. | 15 |
| - **Supplemental Figure S9:** Genomic alterations for from VCaPER differentially expressed proteins with differential TE ratio do not always correlate with altered mRNA expression. | 17 |
| - **Supplemental Figure S10:** NUDT19 expression correlates with determinants of high grade and resistant PCa. | 19 |
| - **Supplemental Figure S11:** lncRNAs are more associated with ribosomes than expected in VCaPER. | 21 |
| - **Supplemental Figure S12:** lncRNAs with high TE ratios in VCaPER are linked to higher PCa grade. | 23 |
| - **Supplemental Figure S13:** Genomic alterations are not always linked to changes in lncRNA expression. | 25 |
| - **Supplemental Figure S14:** Candidate lncRNAs exhibit alternative splicing events in PCa ENZ resistance. | 27 |
| - **Supplemental_Table_S1:** Total and polyribosome RNAseq data - CPM values and EdgeR differential Expression | Supplemental file |
| - **Supplemental_Table_S2:** Protein abundances from mass spectrometry (M.S.) and differential expression analysis | Supplemental file |
| - **Supplemental_Table_S3:** DAVID 6.8 gene ontology term enrichment analysis for Figures 2C et S2A | Supplemental file |
| - **Supplemental_Table_S4:** GSEA 4.1.0. Analysis for TOTAL RNA and RNAs highly bound to ribosomes in VCaPER or VCaPCRPC for Figures 2D and S2B-F | Supplemental file |
| - **Supplemental_Table_S5:** String 11.0 and Cytoscape 3.11.1 GO terms analysis for proteins differentially expressed for Figure 3B,C and S4A-C | Supplemental file |
| - **Supplemental_Table_S6:** GSEA 4.1.0. analysis for whole proteome in VCaPER versus VCaPCRPC for Figure 3B | Supplemental file |
| - **Supplemental_Table_S7:** Translation efficiency analysis for VCaPER and VCaPCRPC | Supplemental file |
| - **Supplemental_Table_S8:** DAVID 6.8 gene ontology term enrichment analysis for Figures 4A,B and S7A | Supplemental file |
| - **Supplemental_Table_S9:** List of putative peptides from non-coding genes in Figure 5B | Supplemental file |

**Supplemental Figure S1**

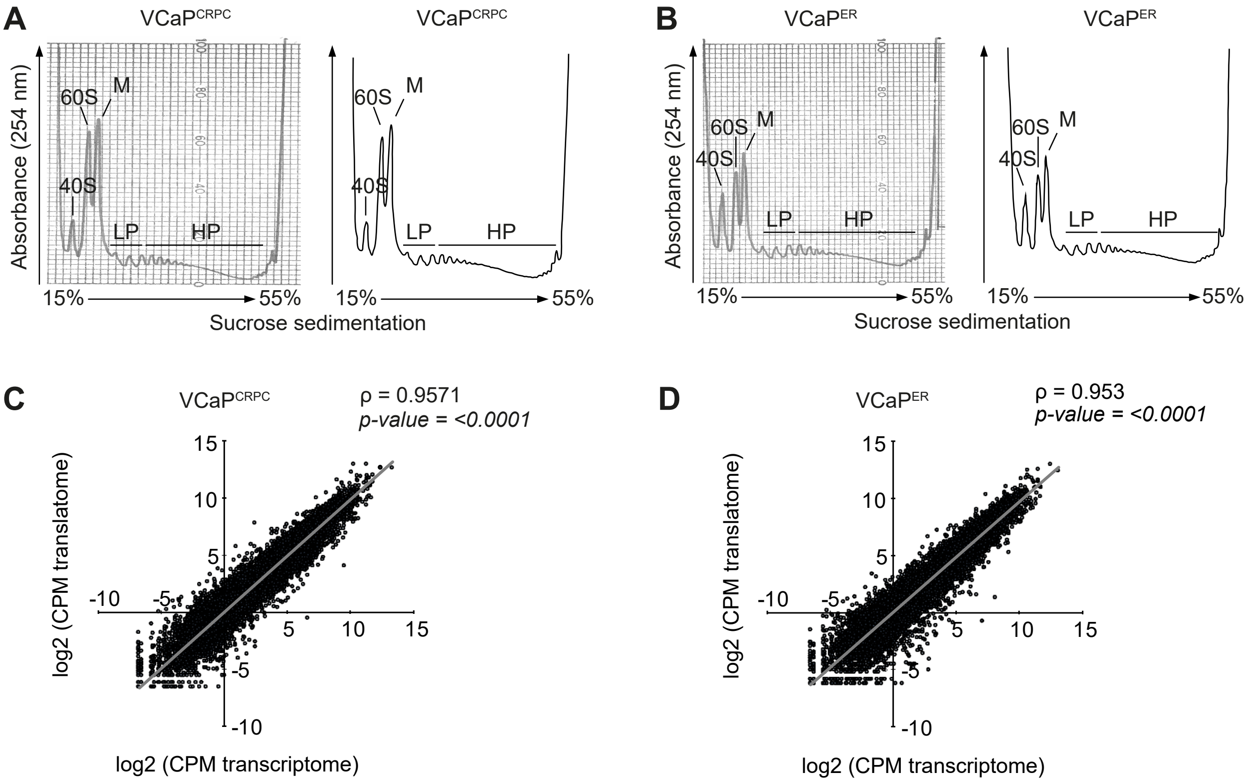


**Figure S1: Transcriptome and translatome analysis of ENZ-sensitive and -resistant cell lines. (A)** Polysome profiling performed in $VCaP^{CRPC}$ and **(B)** $VCaP^{ER}$. Peaks show ribosomal subunits (40S and 60S), mono-ribosomes (M), light (LP) and heavy polysomes (HP), distributed along a sucrose sedimentation gradient. **(C)** Correlation of total mRNA levels with RNA association to ribosomes in $VCaP^{CRPC}$ and **(D)** $VCaP^{ER}$ cell lines. Pearson correlation coefficients (ρ) and linear regression (grey lines) are indicated. CRPC: Castration-Resistant Prostate Cancer, ER: Enzalutamide-Resistant.

### Supplemental Figure S2

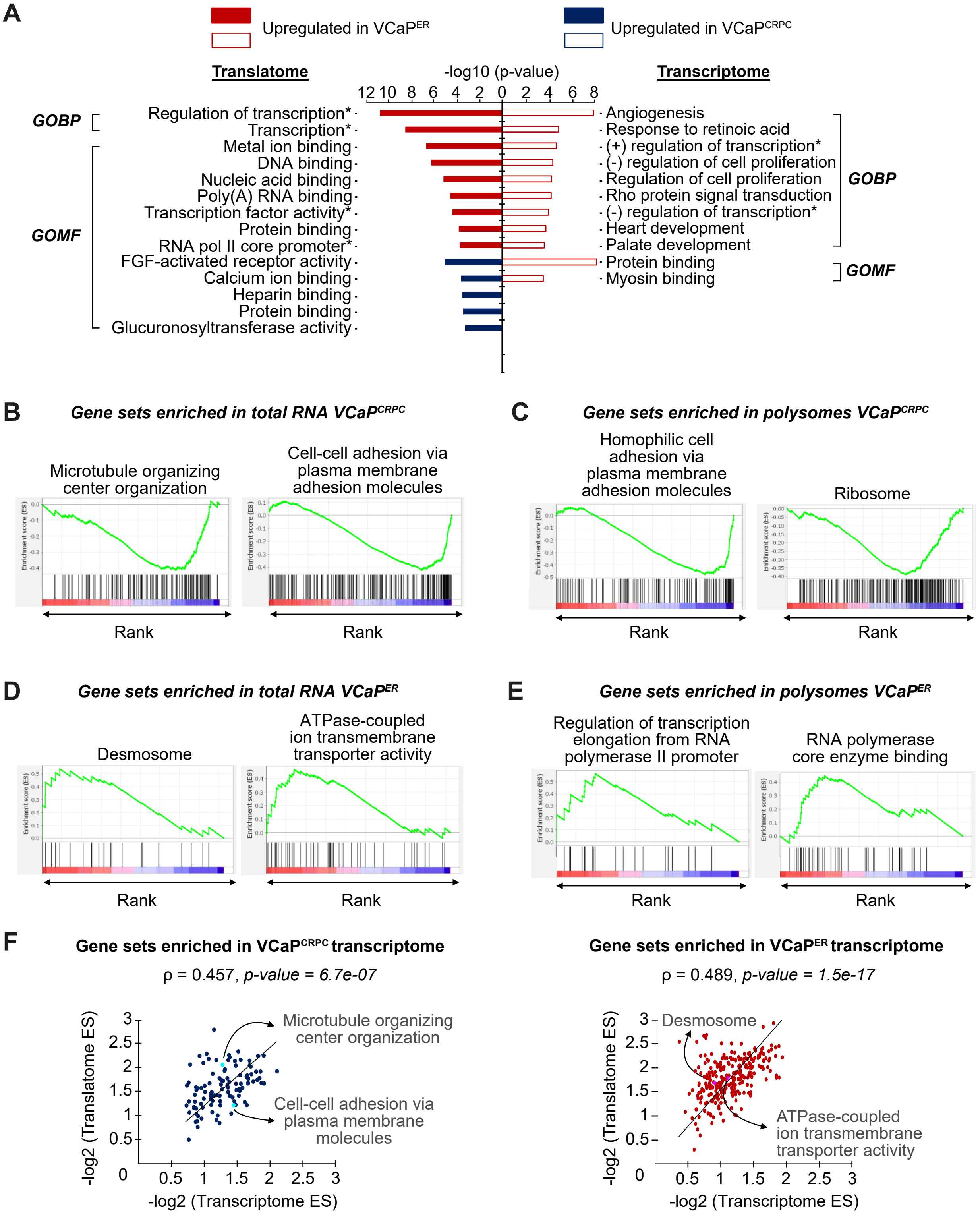

**Figure S2: Significant enrichment of gene sets in VCaP$^{ER}$ and VCaP$^{CRPC}$ transcriptomes and translatomes. (A)** GO-term enrichment analysis showing top 10 significant biological process (GOBP) and molecular function (GOMF) terms enriched in VCaP$^{ER}$ or VCaP$^{CRPC}$. *Full names of GO terms: regulation of transcription - DNA-templated, transcription - DNA-templated, transcription factor activity - sequence-specific DNA binding, RNA polymerase II core promoter proximal region sequence-specific DNA binding, positive regulation of transcription from RNA polymerase II promoter, negative regulation of transcription from RNA polymerase II promoter. **(B)** GSEA enrichment plots for selected gene sets in VCaP$^{CRPC}$ total RNA, **(C)** VCaP$^{CRPC}$ poly-ribosomal fractions, **(D)** VCaP$^{ER}$ total RNA and **(E)** VCaP$^{ER}$ poly-ribosomal fractions. **(F)** Scatterplots of gene set enrichment analysis (GSEA) enrichment scores (ES) for gene sets upregulated in VCaP$^{CRPC}$ (left) and VCaP$^{ER}$ (right) transcriptome. ES scores in VCaP$^{CRPC}$ (blue) or VCaP$^{ER}$ (red) translatomes and transcriptomes are plotted. Spearman correlation coefficients (ρ) and linear regressions (black lines) are indicated.

**Supplemental Figure S3**

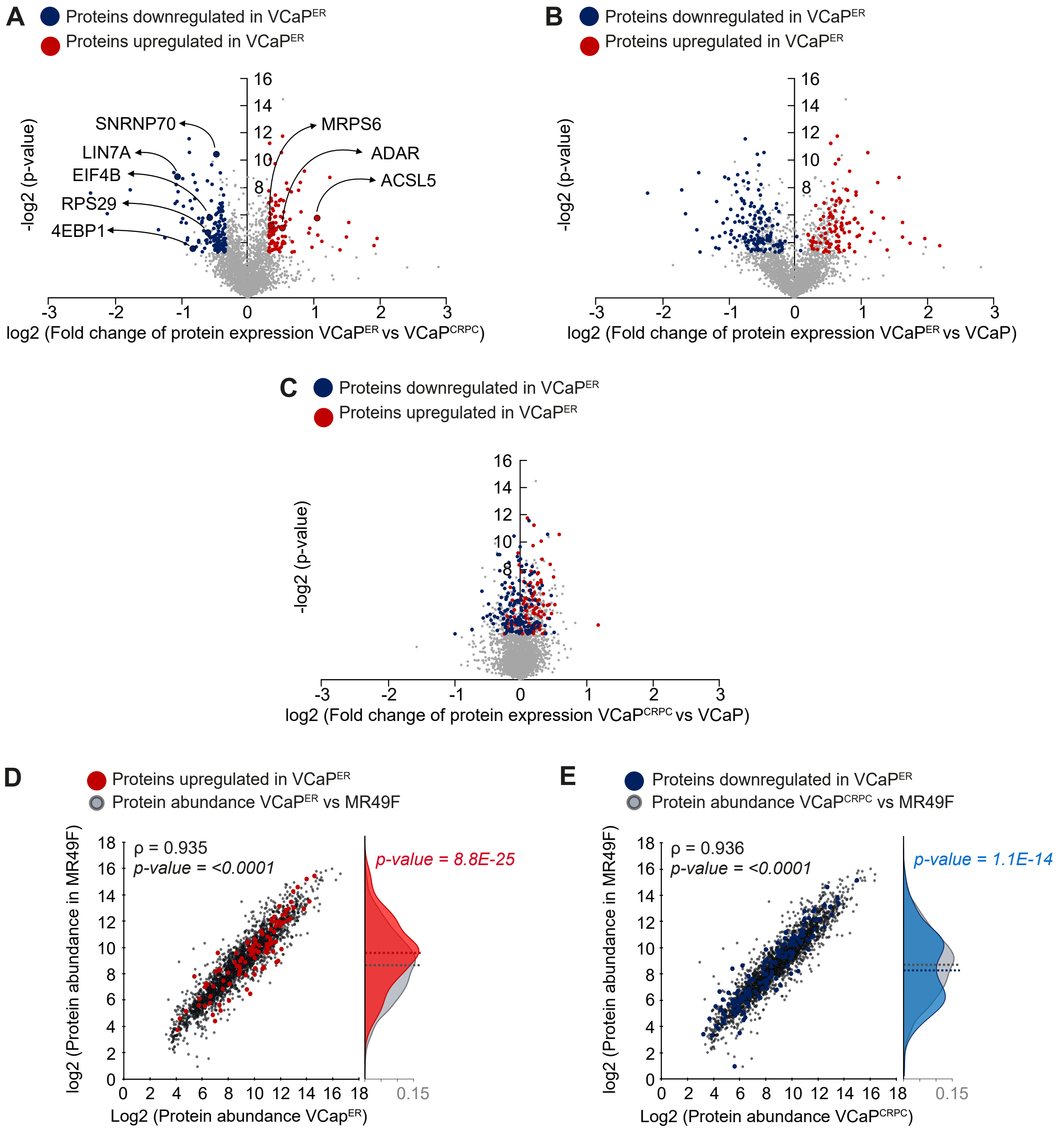

**Figure S3: The VCaP$^{ER}$ ENZ-resistance model recapitulates established features of drug-resistant PCa models. (A)** Volcano plots showing proteins differentially expressed between VCaP$^{ER}$ and VCaP$^{CRPC}$, **(B)** VCaP$^{ER}$ and VCaP, or **(C)** VCaP$^{CRPC}$ and VCaP. Upregulated (red) or downregulated (blue) proteins in VCaP$^{ER}$ compared to VCaP$^{CRPC}$ are indicated. Proteins of interest are highlighted. **(D)** Spearman correlation of protein abundances between ENZ-resistant MR49F and VCaP$^{ER}$ or **(E)** VCaP$^{CRPC}$. Density plots show MR49F expression of proteins **(D)** up or **(E)** downregulated in VCaP$^{ER}$.

**Supplemental Figure S4**

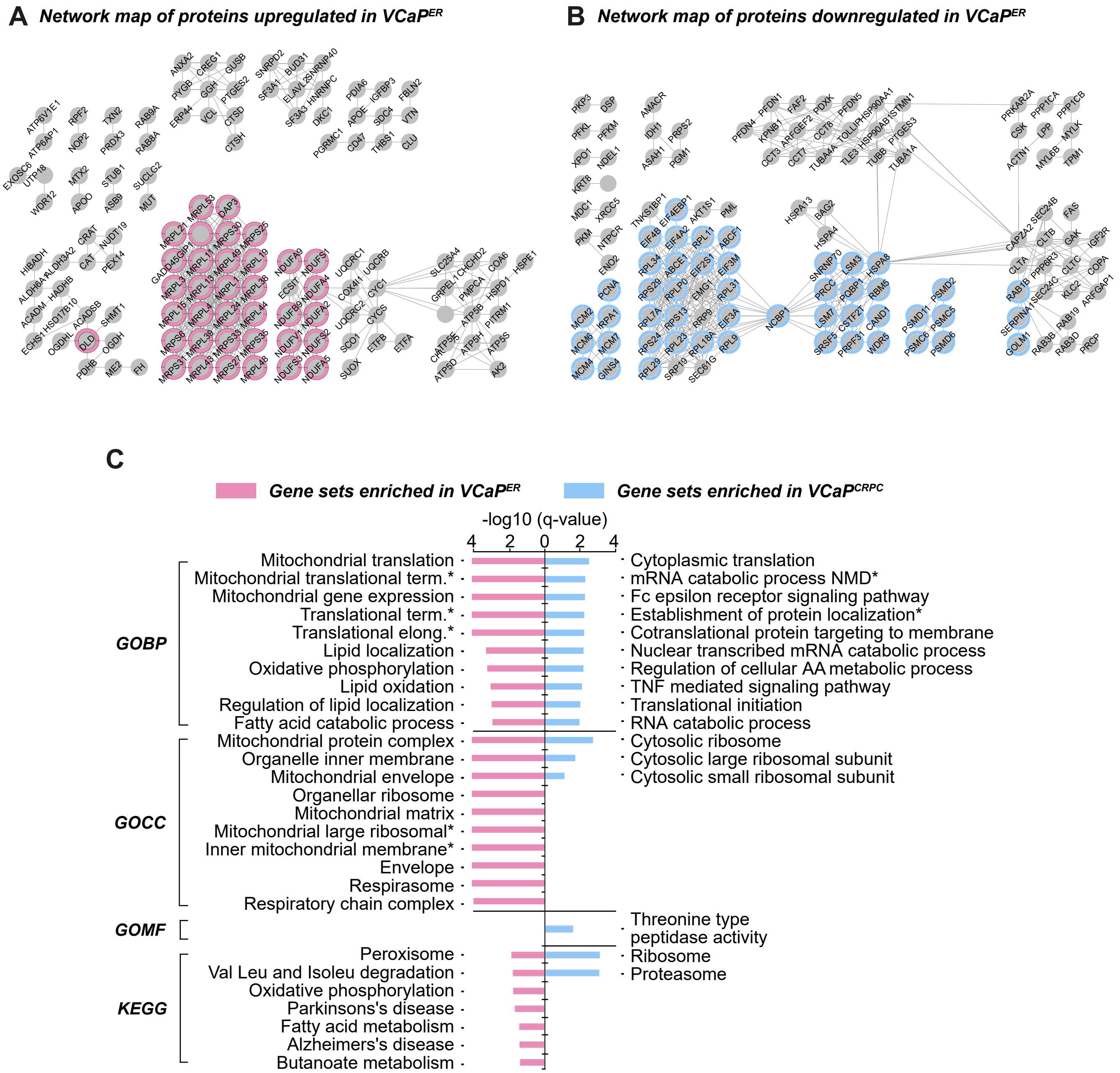

**Figure S4: Proteins up- and downregulated in VCaP$^{ER}$ assemble in protein networks. (A)** Network analysis of proteins differentially expressed between VCaP$^{ER}$ and VCaP$^{CRPC}$, showing protein clusters upregulated in VCaP$^{ER}$ and **(B)** downregulated in VCaP$^{ER}$. **(C)** GSEA shows top 10 enriched gene sets for biological processes (GOBP), cellular components (GOCC), molecular functions (GOMF) and Kyoto Encyclopedia of Genes and Genomes (KEGG) for proteins down (blue) or upregulated (pink) in VCaP$^{ER}$. *Full names of gene sets: mitochondrial translational termination, translational termination, translational elongation, mitochondrial large ribosomal subunit, inner mitochondrial membrane protein complex, nuclear transcribed mRNA catabolic process nonsense mediated decay, establishment of protein localization to endoplasmic reticulum.

### Supplemental Figure S5

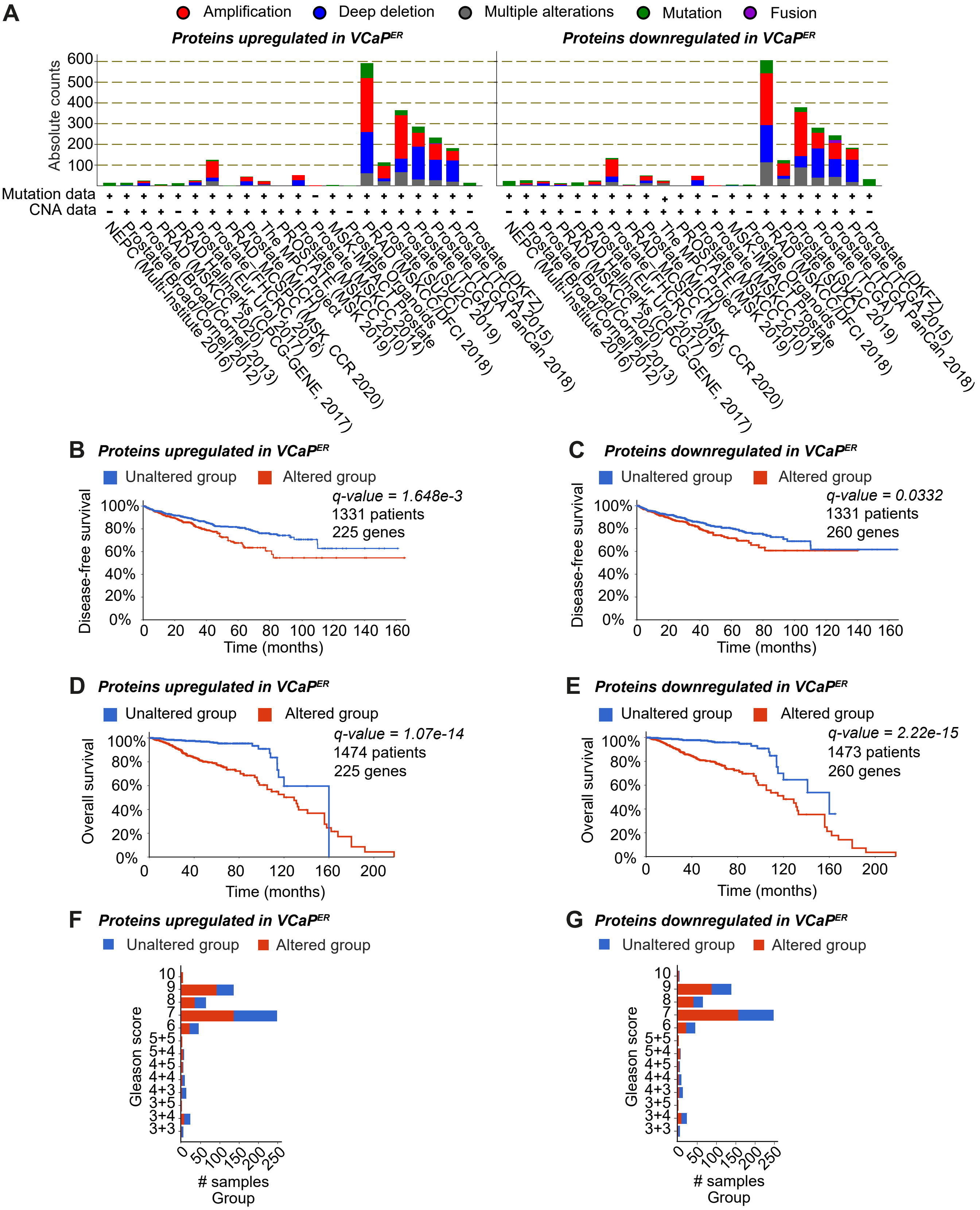

**Figure S5: Genomic alterations in VCaP$^{ER}$-specific proteins are linked to PCa drug resistance in patients. (A)** Gene alteration counts in PCa patient samples for genes coding for proteins up- or downregulated in VCaP$^{ER}$ according to PCa type. **(B)** Kaplan-Meier graph of disease-free PCa patient survival for proteins up- or **(C)** downregulated in VCaP$^{ER}$ and **(D)** overall survival for proteins up- or **(E)** downregulated in VCaP$^{ER}$. **(F)** Distribution of Gleason scores in PCa patients according to genomic alterations in proteins up- or **(G)** downregulated in VCaP$^{ER}$.

**Supplemental Figure S6**

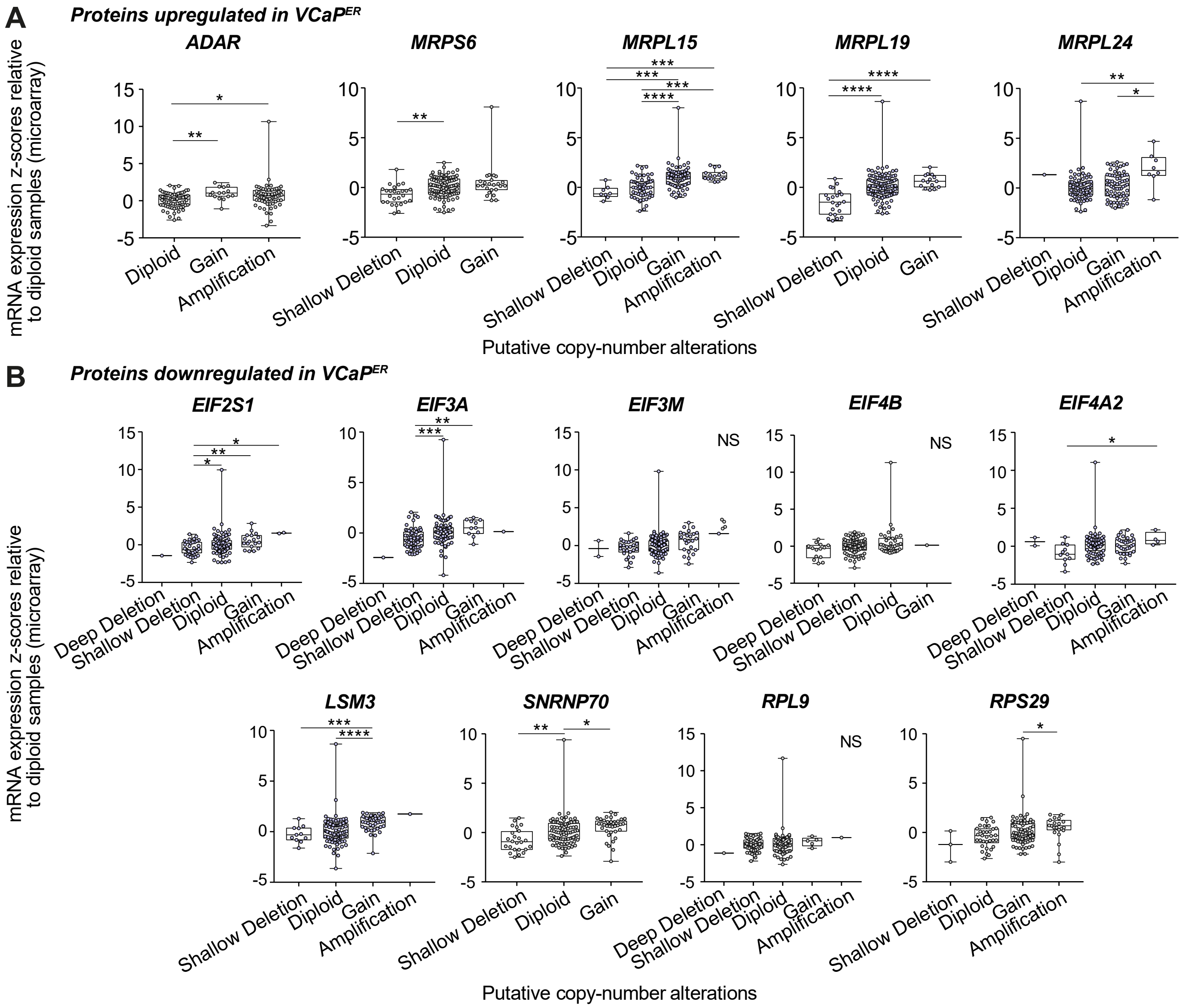


**Figure S6: Genomic alterations are not always linked to RNA expression for genes coding for differentially expressed proteins in VCaP$^{ER}$. (A)** Boxplots showing mRNA expression according to gene alteration type in PCa patient data for proteins up or **(B)** downregulated in VCaP$^{ER}$. Kruskal-wallis test was performed to analyse significant differences in means between groups ( *: $p < 0.05$, **: $p < 0.01$, ***: $p < 0.001$, ****: $p < 0.0001$).

**Supplemental Figure S7**

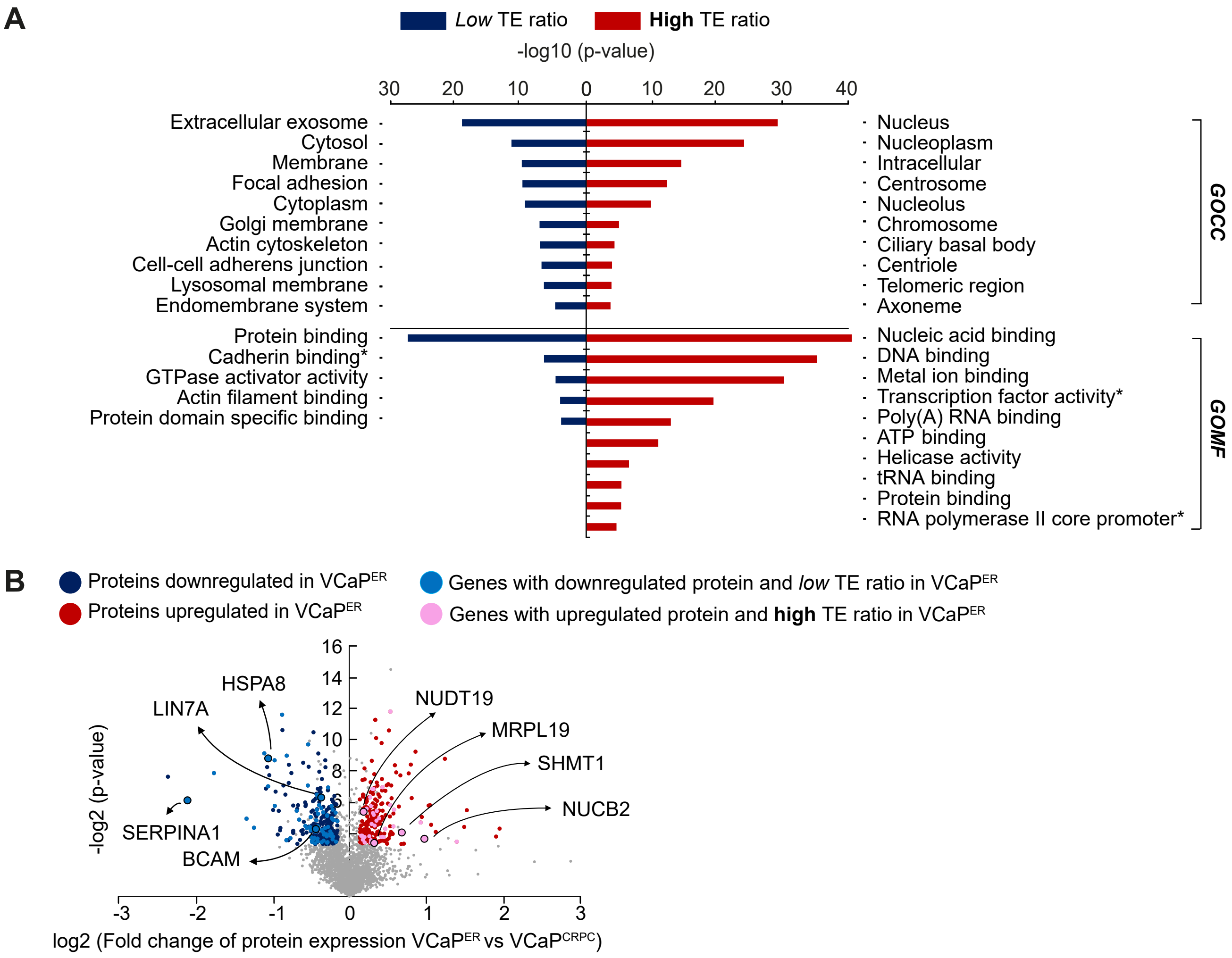


**Figure S7: High TE ratio RNAs recapitulate key GO-terms from the translatome in VCaP$^{ER}$, but do not necessarily lead to higher protein expression. (A)** GO term enrichment analysis shows terms for top 10 significant cellular components (GOCC) and molecular functions (GOMF) enriched in genes with low or high TE ratios in VCaP$^{ER}$ compared to VCaP$^{CRPC}$. *Full names of GO terms: cadherin binding involved in cell-cell adhesion, transcription factor activity, sequence-specific DNA binding, RNA polymerase II core promoter proximal region sequence-specific DNA binding. **(B)** Volcano plot shows proteins differentially expressed between VCaP$^{ER}$ and VCaP$^{CRPC}$. Up or downregulated in VCaP$^{ER}$ are shown in red and blue respectively. Upregulated proteins with significantly upregulated TE in VCaP$^{ER}$ are marked in pink and downregulated proteins with significantly downregulated TE are marked in blue.

**Supplemental Figure S8**

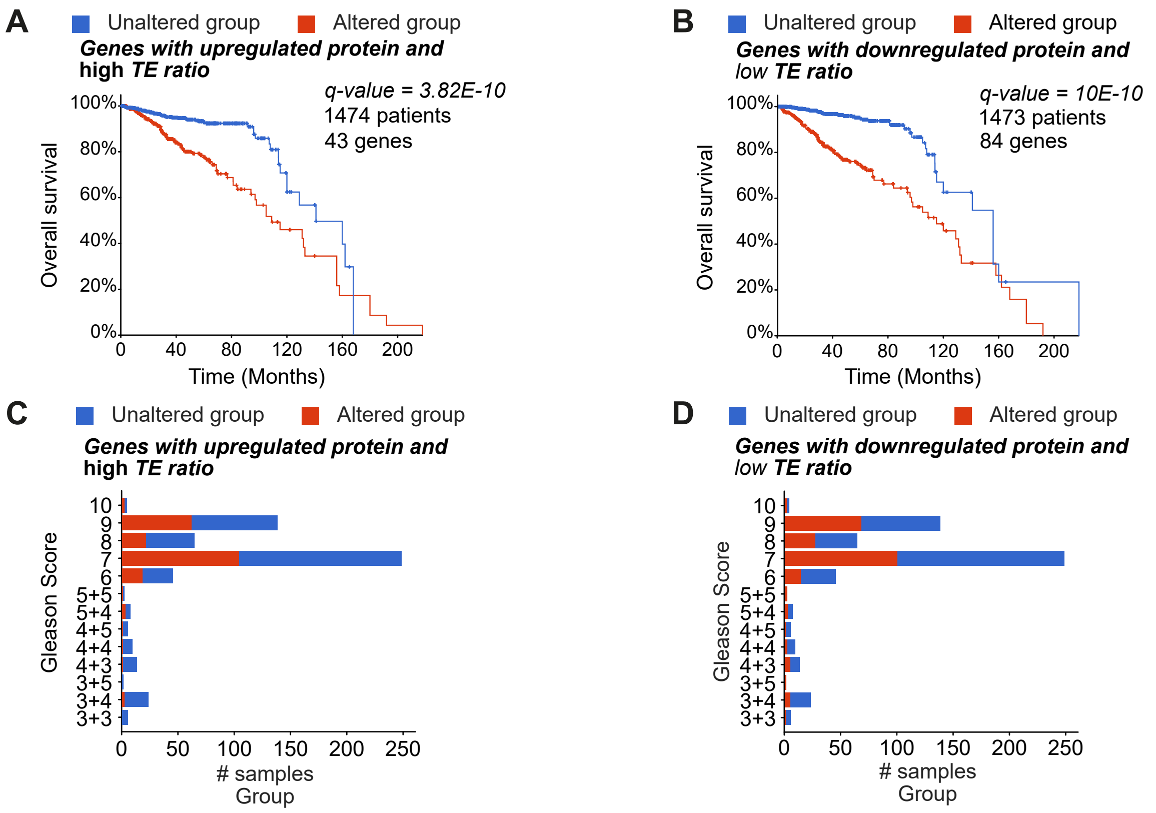


**Figure S8: Genomic alteration in proteins upregulated in VCaP$^{ER}$ is linked to high PCa grade, independently of TE ratio. (A)** Overall survival for PCa patients according to alterations in genes coding for proteins upregulated with high TE ratios in VCaP$^{ER}$ or **(B)** downregulated with low TE ratios. **(C)** Distribution of Gleason scores for patients according to alterations in genes coding for proteins upregulated with high TE ratios or **(D)** downregulated with low TE ratios.

### Supplemental Figure S9

**A** ***Genes with upregulated proteins and*** **high** ***TE ratio***

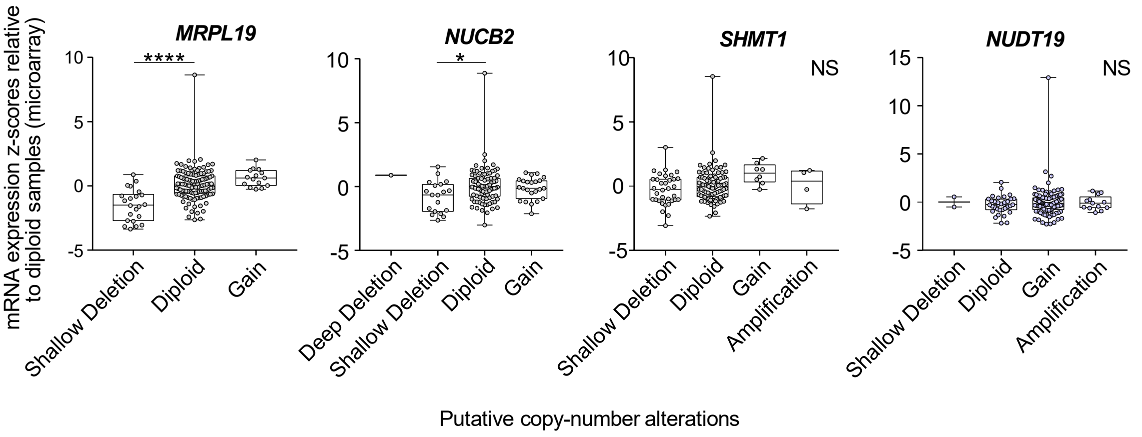


**B** ***Genes with downregulated proteins and*** *low* ***TE ratio***

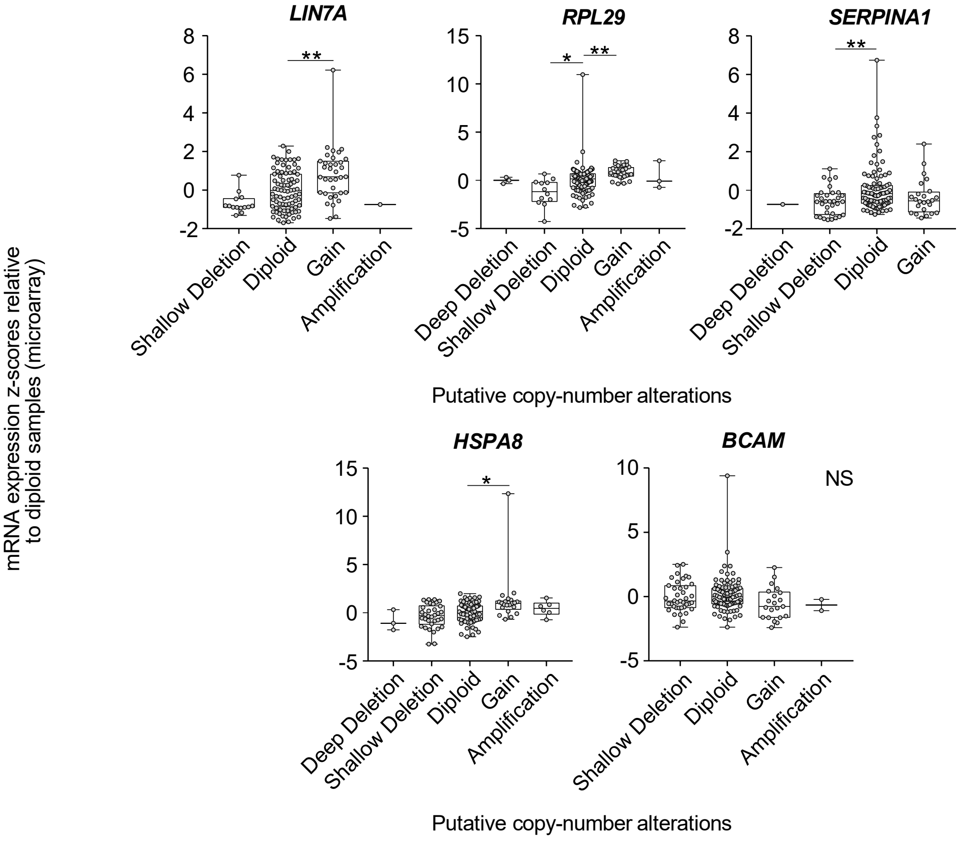

**Figure S9: Genomic alterations for from VCaP$^{ER}$ differentially expressed proteins with differential TE ratio do not always correlate with altered mRNA expression. (A)** mRNA expression according to type of genomic alteration in PCa patients, for genes coding for proteins upregulated and with high TE ratios in VCaP$^{ER}$ or **(C)** downregulated with low TE ratios. Significance was assessed with Kruskal-wallis test ( *: $p < 0.05$, **: $p < 0.01$, ***: $p < 0.001$, ****: $p < 0.0001$).

**Supplemental Figure S10**

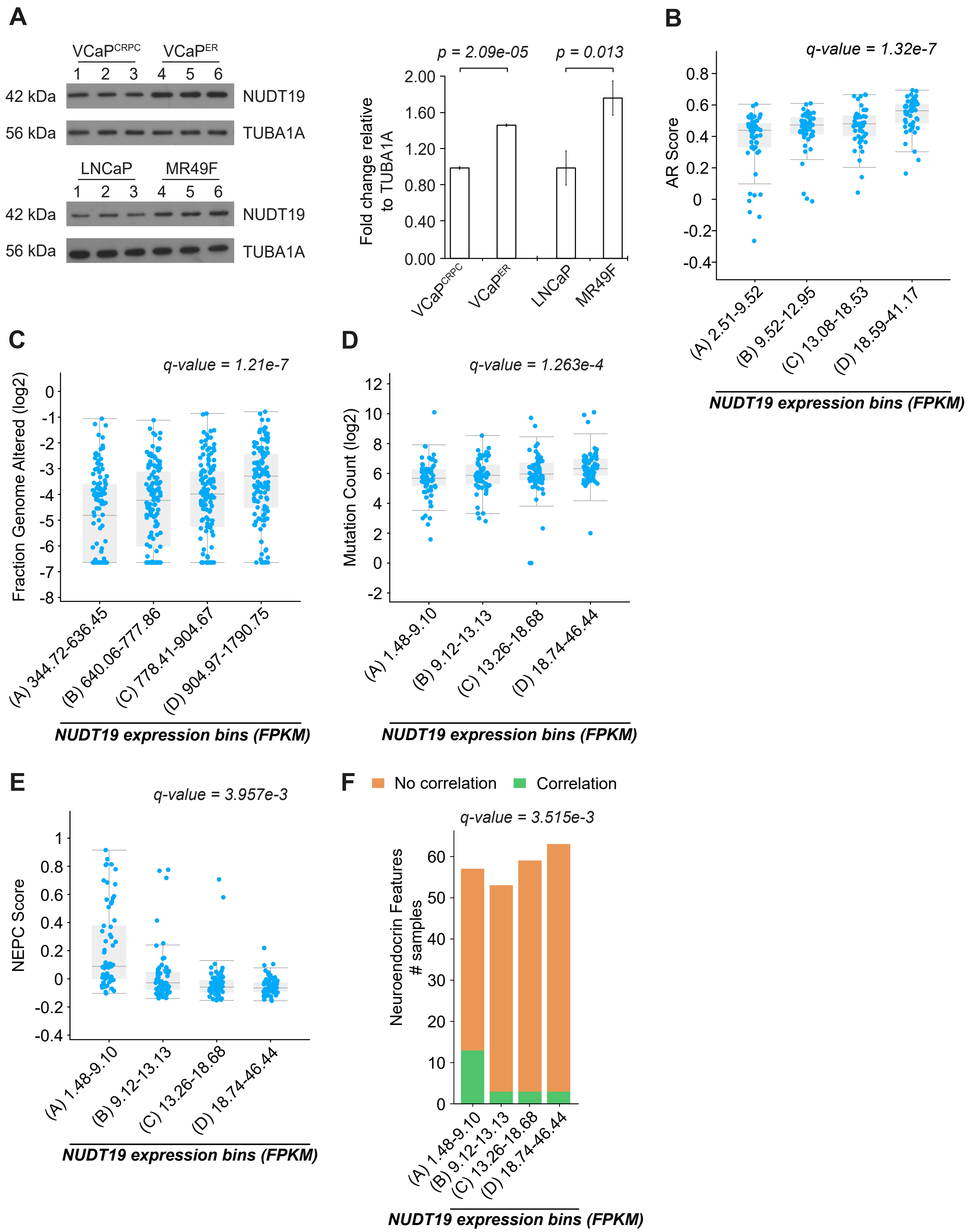

**Figure S10: *NUDT19* expression correlates with determinants of high grade and resistant PCa. (A)** Protein expression of NUDT19 in $VCaP^{ER}$, $VCaP^{CRPC}$, LNCaP and MR49F, shown in triplicate lanes (left). Quantification of NUDT19 protein levels in $VCaP^{ER}$ or MR49F respectively normalized to $VCaP^{CRPC}$ or LNCaP (right). **(B)** AR score, **(C)** fraction of the genome altered, **(D)** mutation counts, **(E)** NEPC score and **(F)** number of PCa patient samples containing neuroendocrine features according to *NUTD19* expression binning in PCa patients.

### Supplemental Figure S11

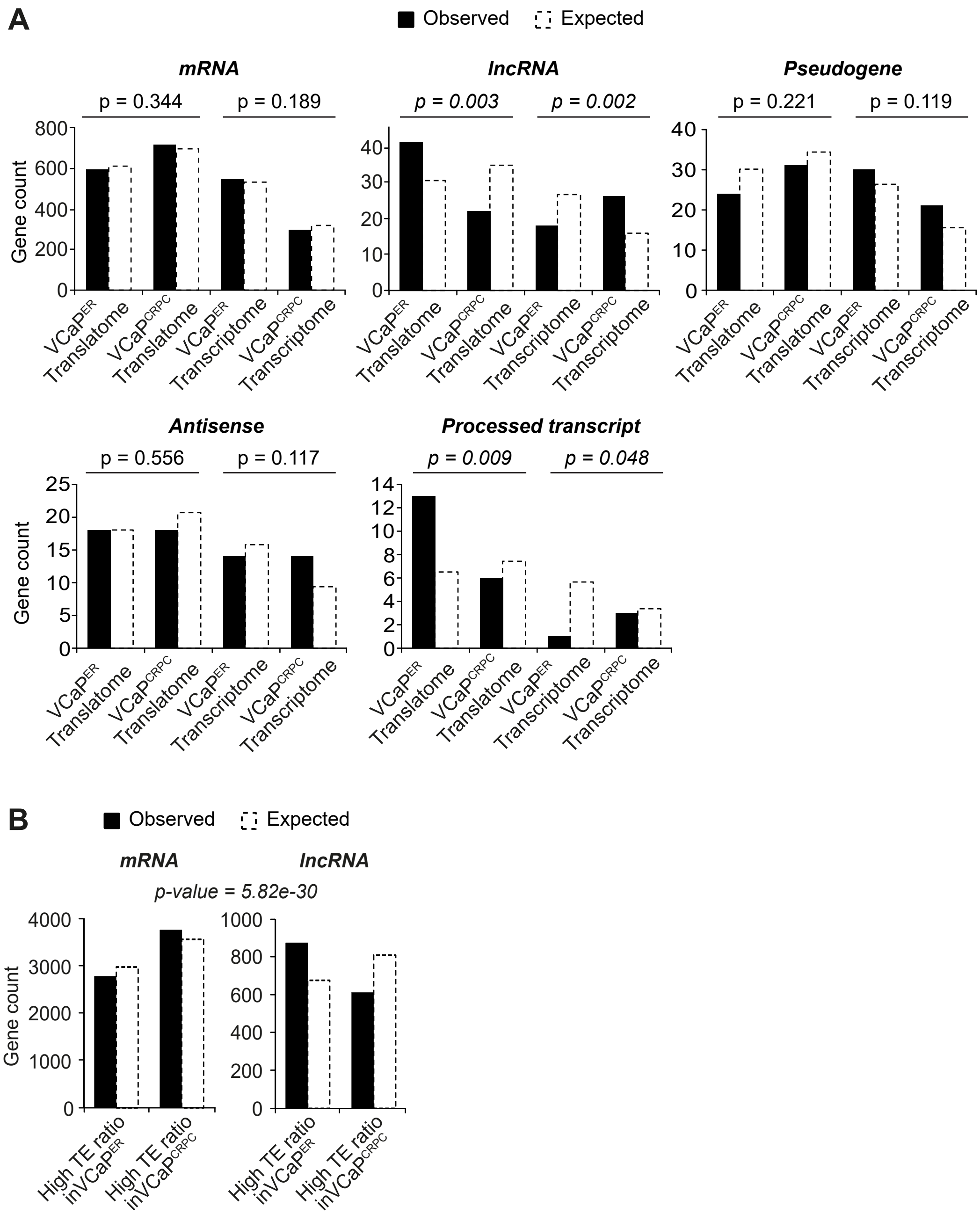


**Figure S11: lncRNAs are more associated with ribosomes than expected in VCaP$^{ER}$. (A)** Observed and expected gene counts for different categories of genes from both the translatome and transcriptome in VCaP$^{ER}$ compared to VCaP$^{CRPC}$. **(B)** Observed and expected gene counts of coding genes and lncRNAs with high translation efficiencies in VCaP$^{ER}$ compared to VCaP$^{CRPC}$.

### Supplemental Figure S12

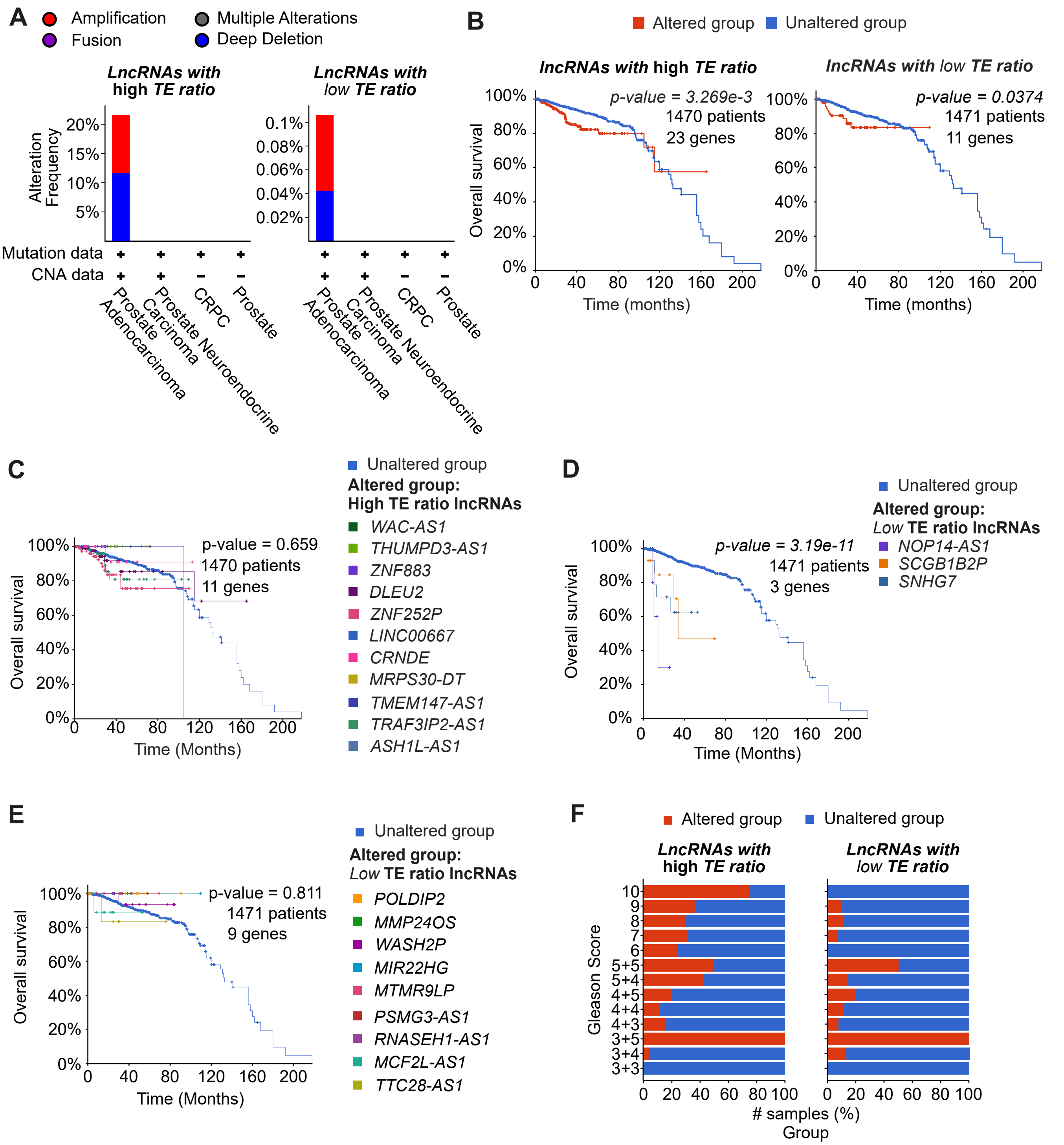

**Figure S12: lncRNAs with high TE ratios in VCaP$^{ER}$ are linked to higher PCa grade. (A)** Alteration frequencies for lncRNAs with high or low TE ratios in VCaP$^{ER}$ compared to VCaP$^{CRPC}$ according to PCa type. CNA : copy-number alteration. **(B)** Overall survival for PCa patients with or without alterations in lncRNAs with high (left) or low (right) TE ratios in VCaP$^{ER}$. **(C)** Overall survival for PCa patients with alterations in single lncRNAs with high TE ratios that show no significant link to patient survival, **(D)** lncRNAs with low TE that are correlated with lower patient survival, or **(E)** lncRNAs with low TE ratios that show no significant link to patient survival. **(F)** Percentage of PCa patient samples per Gleason score with high (left) or low (right) TE lncRNAs with or without alterations.

**Supplemental Figure S13**

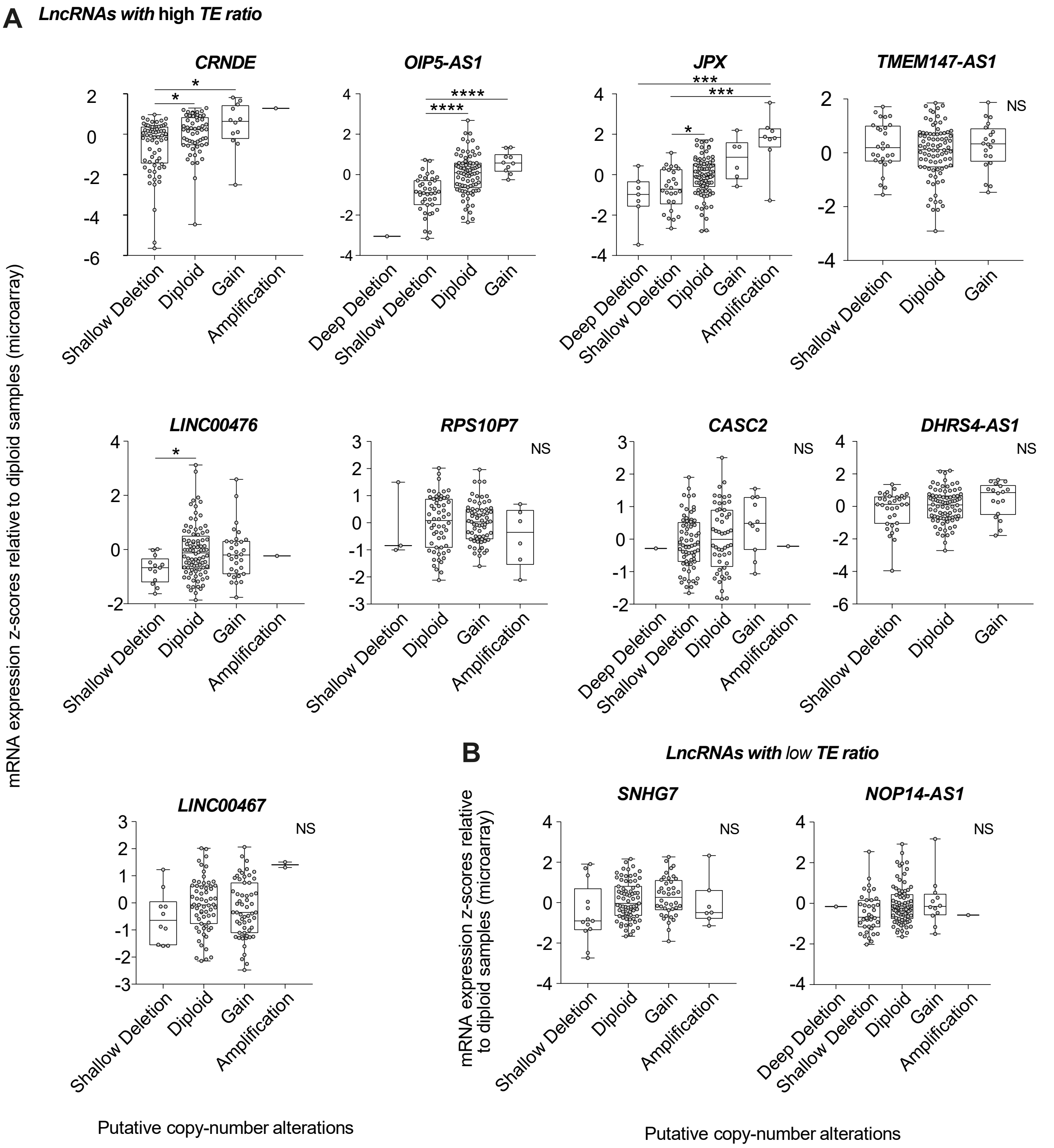


**Figure S13: Genomic alterations are not always linked to changes in lncRNA expression. (A)** RNA expression according to type of genomic alteration in PCa patients for lncRNAs with high TE ratios in VCaP$^{ER}$ or **(B)** low TE ratios in VCaP$^{ER}$. Significance was assessed with Kruskal-wallis test ( *: $p < 0.05$, **: $p < 0.01$, ***: $p < 0.001$, ****: $p < 0.0001$).

### Supplemental Figure S14

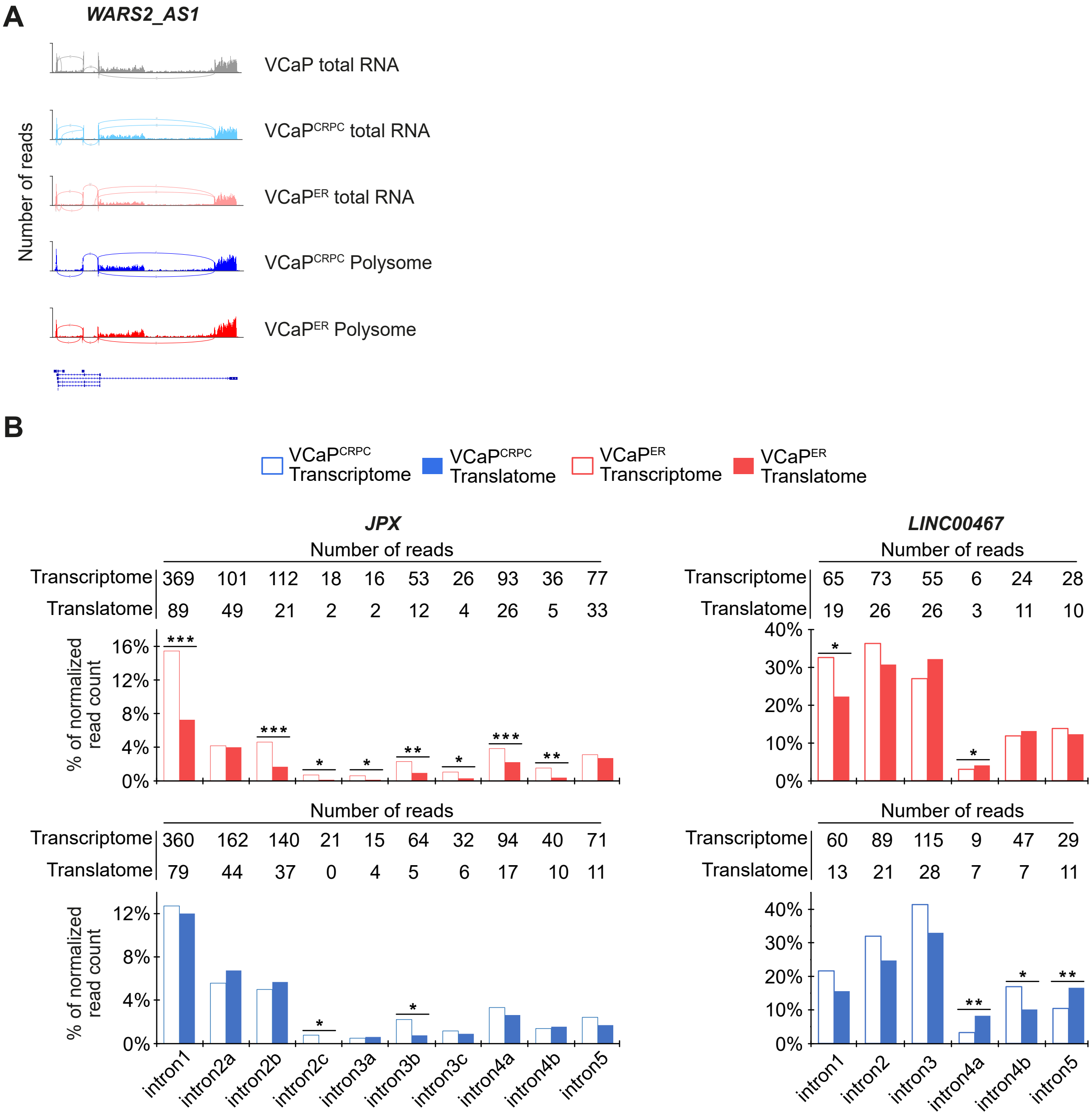


**Figure S14: Candidate lncRNAs exhibit alternative splicing events in PCa ENZ resistance. (A)** Sashimi plots for the lncRNA WARS2_AS1 in PCa cell lines shows alternatively spliced isoforms in the transcriptome and translatome of VCaP$^{ER}$ and VCaP$^{CRPC}$. **(B)** Quantification of *JPX* and *LINC00467* lncRNAs split reads count, corresponding to all splicing events in the transcriptome and translatome of VCaP$^{CRPC}$ (blue) or VCaP$^{ER}$ (red) ( *: $p < 0.05$, **: $p < 0.01$, ***: $p < 0.001$).
